# Anti-GABA_B_ receptor autoimmunity causes cognitive decline by presynaptic pathology

**DOI:** 10.64898/2026.10.22.761036

**Authors:** Josefine Sell, Eleonora Anna Loi, Vahid Rahmati, Myrtill Majoros, Toni Kirmann, Holger Haselmann, Alexander Stumpf, Torsten Götz, Christiana Franke, Frank Leypoldt, Sven Jarius, Nico Melzer, Jan Lewerenz, Albrecht Kunze, Harald Prüss, Dietmar Schmitz, Stefan Hallermann, Christoph Schmidt-Hieber, Hans-Christian Kornau, Christian Geis

## Abstract

The GABA_B_ receptor is a central regulator of neuronal transmission and excitability by pre- and postsynaptic mechanisms. Its dysfunction causes severe neurological disease, but the underlying pathomechanisms remain elusive. Here, we exploit a distinct CNS autoimmune disorder in which patients develop specific autoantibodies against GABA_B_ receptors causing autoimmune encephalitis with severe memory dysfunction and epilepsy. We demonstrate that human autoantibodies target the sushi-domains of the GABA_B_ receptor B1a subunit and the B2 subunit and interfere with presynaptic signaling. This specific presynaptic pathology results from exclusive functional antagonism of the B1a subunit and disrupts short- and long-term synaptic plasticity, abnormally amplifies hippocampal network oscillations and coupling *in vivo*, and ultimately impairs memory consolidation. Our results demonstrate that autoimmune presynaptic pathomechanisms alone are sufficient to interfere with memory processing, thus defining a causal link between molecular GABA_B_ receptor disturbance, synaptic pathology, circuit dysfunction, and dementia distinct from classical neurodegenerative processes.

## Introduction

GABA_B_ receptors (GABA_B_Rs) are class C family G-protein-coupled receptors (GPCRs) that are crucial in modulating synaptic transmission over timescales from seconds to minutes^1^. As critical regulators of neuronal excitability, synaptic transmission, and neural circuits dynamics, altered GABA_B_R signaling have been implicated in the pathophysiology of various central nervous system (CNS) disorders, including neurocognitive disorders, epilepsy, schizophrenia, and autism spectrum disorders^1–6^. Pathogenic GABA_B_R subunit variants cause severe neurodevelopmental delay, intellectual disability, and epilepsy in human^4,7,8^. Consistently, GABA_B1_R subunit knockout mice exhibit neurodevelopmental defects, epileptiform activity, and memory dysfunction^9–11^.

Strikingly, GABA_B_Rs have also been identified as targets of highly specific pathogenic autoantibodies (aAbs) in serum and cerebrospinal fluid (CSF) of patients with an autoimmune CNS disorder, defining anti-GABA_B_R autoimmune encephalitis (AE)^12^. This condition is characterized by severe limbic encephalitis, presenting with pronounced anterograde memory impairment, confusion, and temporal lobe seizures^13^. Early immunotherapeutic interventions, such as plasma exchange to remove aAbs or B-cell depletion with rituximab, improve symptoms, underscoring the pathogenic role of these aAbs^14,15^. Importantly, this acute autoimmune condition with highly specific aAbs offers a unique opportunity to understand direct and immediate GABA_B_R-mediated pathology *in vivo* and its consequences for neuronal and brain circuit function in a clinically relevant context.

GABA_B_Rs are obligate heterodimers composed of a GABA_B1_ subunit (either GABA_B1a_ or GABA_B1b_) and a GABA_B2_ subunit^1,16^. The subunits’ architecture features a large extracellular Venus flytrap domain and a heptahelical transmembrane domain. While only the GABA_B1_ subunit contains the ligand-binding site for GABA and synthetic agonists like baclofen (β-p-Cl-phenyl-γ-amino-butyric acid) and phenibut (4-amino-3-phenyl-butyric acid)^17,18^, the intracellular loops of GABA_B2_ mediate the G-protein coupling^1^. The two GABA_B1_ isoforms (GABA_B1a_ and GABA_B1b_) arise from alternative transcription initiation sites, conferring a pair of sushi-domains (SDs) specifically at the N-terminus of GABA_B1a_. These SDs act as an axonal targeting signal, directing GABA_B1a_-containing receptors to presynaptic terminals^11,19,20^. These presynaptic GABA_B_Rs localize mainly perisynaptically, functioning as autoreceptors at inhibitory synapses by controlling GABA release and as heteroreceptors at excitatory synapses by inhibiting glutamate release^21,22^ (**Fig. 1a**). Mechanistically, this transmitter release reduction arises from the inhibition of voltage-gated Ca^2+^ channels via G-protein signaling and/or direct interference with the vesicle release machinery^23,24^. Conversely, GABA_B1b_-containing receptors localize postsynaptically, where they activate inwardly-rectifying potassium channels (GIRKs/Kir3), resulting in hyperpolarization of postsynaptic membranes and reduced Ca^2+^ influx^16,25^.

**Fig 1:**
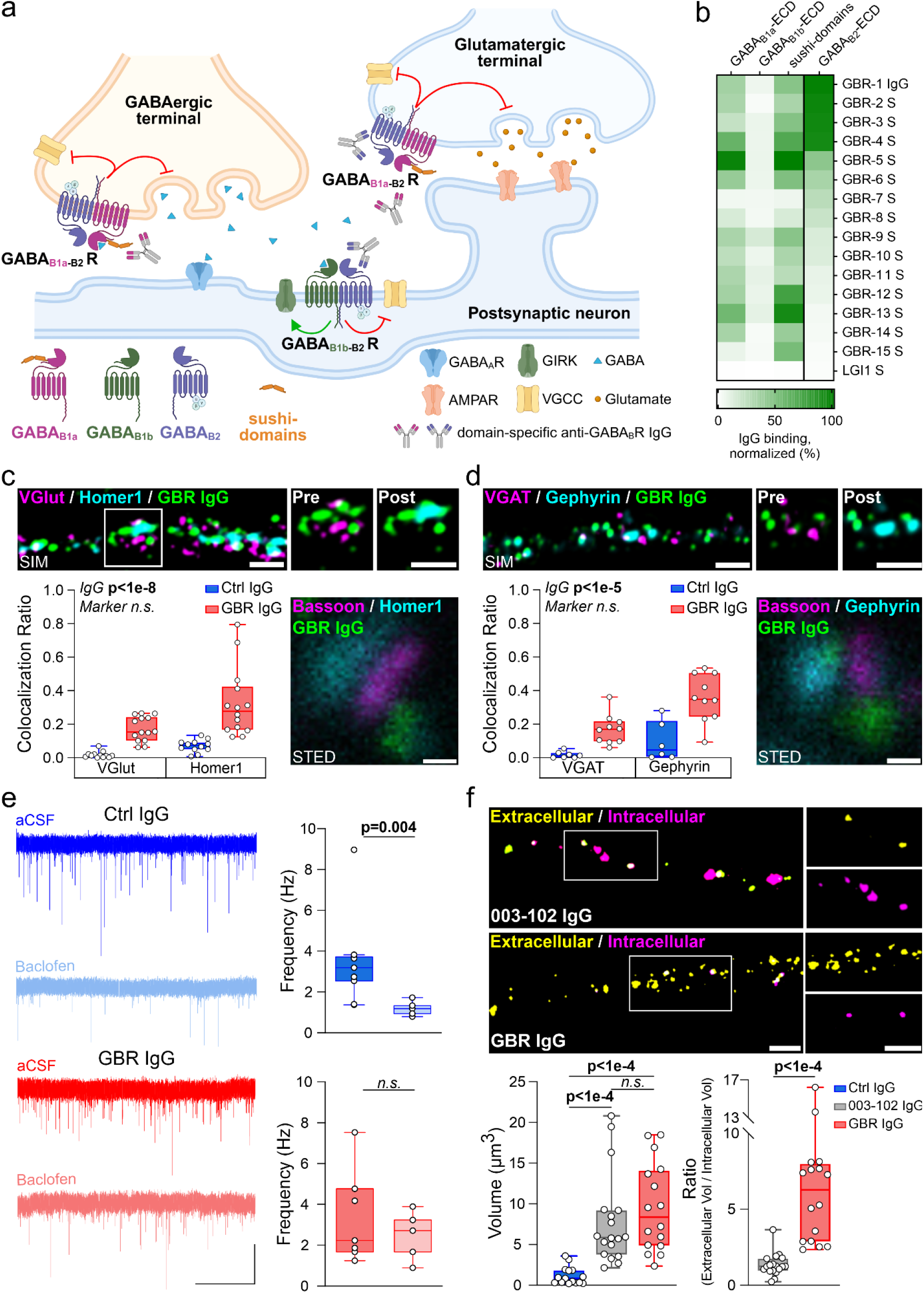
**a**) Schematic illustrating GABA_B_R synaptic distribution and subunit compositions. **b**) IgG binding of patients’ sera (GBR S, 1:100) and purified IgG (GBR-1 IgG, 1:1000) to soluble Fc fusion proteins encompassing the extracellular domains (ECDs) of the GABA_B_R subunits GABA_B1a_, GABA_B1b_, GABA_B2_ or isolated sushi-domains determined by ELISA. mGluR5-ECD-Fc served as control antigen. LGI1 S: control serum of a patient with IgG aAbs to the synaptic protein LGI1. Green shading indicates percentage of IgG binding relative to GBR-1 IgG binding to GABA_B2_-ECD. **c**) GBR IgG localization relative to glutamatergic pre- and postsynaptic compartments in primary neurons. Top row: example lattice SIM images (scale bar: 1 µm) of GBR IgG overlaid with VGlut (pre) or Homer1 (post). Bottom left: colocalization ratio (Ctrl IgG vs. GBR IgG: n=12 vs. 14; Linear Mixed-Effects Model [LMM] main effects analysis). Bottom right: example STED image showing perisynaptic localization of GBR IgG immunoreactivity in excitatory synapses (Bassoon: presynaptic active zone; Homer1 postsynaptic density; scale bar: 100 nm). **d**) Same as (C), but for GABAergic pre- and postsynaptic compartments. Example lattice SIM images show GBR IgG overlaid with VGAT (pre) or Gephyrin (post). Colocalization ratio: n=6 vs. 10. **e**) Example recordings (left) and analysis (right) of spontaneous EPSC in primary neurons after 24 h incubation with GBR IgG in naïve aCSF and after baclofen perfusion (scale bars: 5s, 5pA; Ctrl IgG/aCSF: n=6; Ctrl IgG/baclofen: n=5; GBR IgG/aCSF: n=7; GBR IgG/baclofen: n=5; Mann-Whitney U-test). **f**) Top: example confocal images of extracellular (yellow) and intracellular (magent**a**) localization of receptor-bound IgG (scale bar: 2 µ**m**) in dendrites of primary neurons. 003-102 IgG: monoclonal anti-GluN1 aAb (NMDAR) as positive control for internalization assay. Bottom-left: intravital binding (immunoreactive volume) of 003-102 IgG (n=18) and GBR IgG (n=16) but not Ctrl IgG (n=15) to dendritic surface (Kruskal-Wallis test with Dunn’s post-hoc tests). Bottom-right: extra-vs. intracellular IgG localization ratios after 24 h incubation (Mann-Whitney U-test). Boxplots show median and interquartile range. *n.s.*, not significant. Statistical details in **Extended Data Table 2.** See also **Extended Data** Figure 1a-b.

Understanding how molecular perturbations at the level of individual synapses give rise to complex brain dysfunctions, such as cognitive decline and dementia remains a central challenge in neuroscience. Here, integrating affinity-purified IgG from patients with anti-GABA_B_R AE with multimodal analyses spanning *in vitro* and *in vivo* mouse models, super-resolution imaging, behavioral analyses, electrophysiology, and computational modeling, we define the pathophysiological cascade through which anti-GABA_B_R aAbs induce presynaptic pathology. Our findings identify immune-mediated synaptic dysfunction as a sufficient driver of a persistently hyperactive and inflexible hippocampal state that compromises memory consolidation.

## Results

### Autoantibodies to the GABA_B_R react with GABA_B1a_ and GABA_B2_ subunit epitopes and bind to pre- and postsynaptic terminals

We collected serum samples from 15 patients with autoimmune encephalitis who scored positive for anti-GABA_B_R aAbs in cell-based assays (for details see **Extended Data Table 1**). To determine the epitope reactivity, we tested their binding to four recombinantly expressed extracellular domains (ECDs) of the GABA_B_R using an ELISA-based approach (**Fig. 1b**). The sera showed binding to GABA_B1a_ and to the isolated SDs, which was the main serum reactivity in nearly half of the patients (7/15), but almost no reactivity to GABA_B1b_. Interestingly, several of the sera, predominantly serum samples of patients with associated tumors, showed an additional strong reactivity with the GABA_B2_-ECD (**Fig. 1b, Extended Data Table 1**).

Patient-derived purified IgG and serum samples, but not the IgG fraction of a healthy human donor (Ctrl IgG), bound to vital primary murine hippocampal neurons (14 DIV) and colocalized with a custom-made antibody against the GABA_B1_ subunit^26^ and a commercial antibody against the GABA_B2_ subunit (**Extended Data Fig. 1a**), reflecting the expected distribution of GABA_B_Rs^11,21^. To determine the sample reactivity to neuronal subcellular compartments, we chose the purified IgG with strong reactivity to both GABA_B1a_ and GABA_B2_ subunits (GBR-1 IgG, see **Fig. 1b**; hereinafter referred to as GBR IgG) and performed *super-resolution* lattice-SIM and STED imaging after preincubation of primary neurons. Consistent with the reactivity pattern identified in the ELISA assay, GBR IgG showed pre- and postsynaptic binding at perisynaptic sites of both excitatory and inhibitory synapses indicative of binding to GABA_B_ auto- and heteroreceptors at presynaptic boutons, as well as to postsynaptic GABA_B_Rs (**Fig. 1c-d; Extended Data Fig. 1b**).

To test the functional effects of GBR IgG, we recorded spontaneous excitatory postsynaptic currents (sEPSC) in primary neurons (DIV 11-14). While bath perfusion of the GABA_B_R agonist baclofen reduced sEPSC frequency in Ctrl IgG-treated cells, this effect was abolished in cells pretreated with the GBR IgG (**Fig. 1e**). This implies either an aAb-induced loss of GABA_B_R surface expression or a functional interference of GBR IgG with GABA_B_R agonist activation. As receptor crosslinking and internalization is a dominant pathomechanism in other aAb-mediated AE (e.g. via anti-NMDA receptor aAbs^27,28^), we quantified GABA_B_R synaptic surface and intracellular expression in primary neurons. However, 24 h incubation with GBR IgG did not result in relevant loss of surface receptor expression, consistent with previous observations^29^. Conversely, the positive control using monoclonal 003-102 IgG directed against the NMDA receptor (NMDAR)^30^ caused substantial receptor internalization (**Fig. 1f**).

In summary, these findings demonstrate that the GABA_B1a_ SDs and the GABA_B2_ subunit represent dominant aAb epitopes in anti-GABA_B_R encephalitis. Patient-derived IgG reactive to both extracellular domains of the GABA_B_R binds to pre- and postsynaptic terminals and reduces agonist-induced GABA_B_R activation independently of receptor internalization.

### GABA_B_R IgG induce memory dysfunction in an intraventricular passive-transfer animal model

Next, to model a prominent disease phenotype, we investigated *in vivo* effects of GBR IgG on memory function in mice during continuous 14-day intraventricular IgG infusion via osmotic pumps^31^ (**Fig. 2a**). For this purpose, novel object recognition (NOR) and location (NOL) tests with different retention times were performed to evaluate initial memory encoding and memory consolidation. Three hours after the familiarization phase, both Ctrl and GBR IgG-infused mice groups exhibited equivalent preference for exploring the novel object in NOR (**Fig. 2b**) and NOL (**Extended Data Fig. 2a**) tests, demonstrating unaltered initial memory encoding. However, 24 hours after familiarization, the ability of GBR IgG-treated mice to discriminate between novel and familiar objects was impaired (**Fig. 2c**). This specific deficit in long-term memory consolidation, but not initial acquisition, phenocopies GABA_B1a_-knockout mouse models^11,32^. This suggests a potential subunit-specific antagonistic effect of GBR IgG with the GABA_B1a_ subunit.

**Fig. 2:**
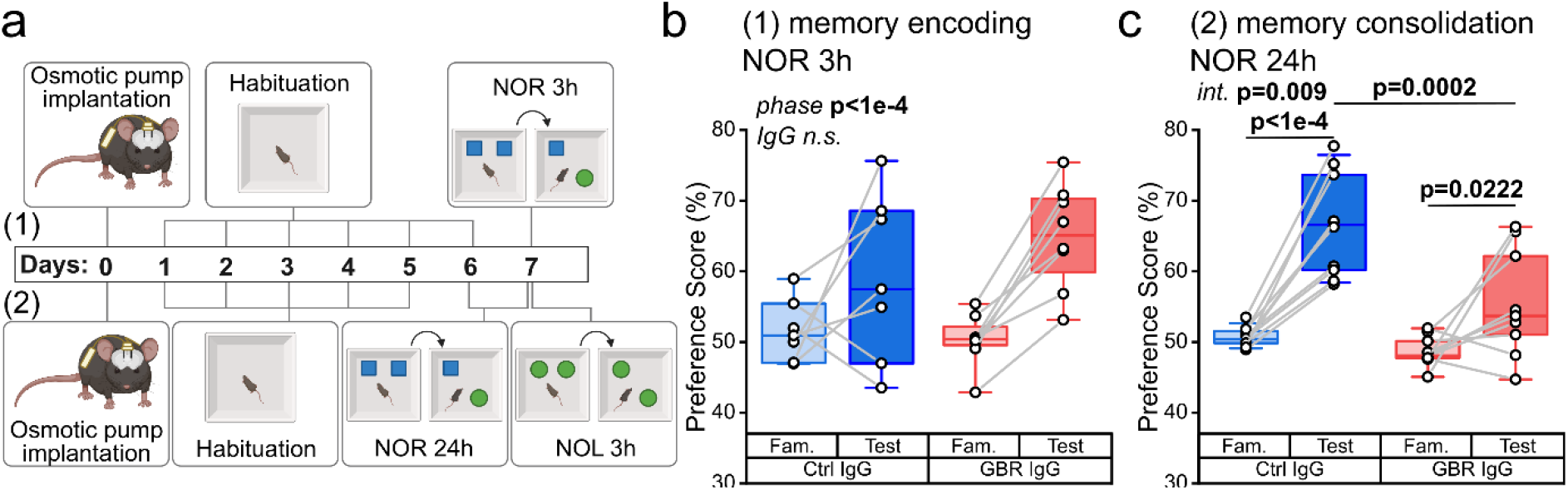
GBR IgG impairs memory consolidation upon intraventricular passive transfer. **a)** Experimental time course of novel object recognition (NOR) and location (NOL) tests to evaluate memory encoding (3 h retention time) or memory consolidation (24 h retention time). **b)** Mice after Ctrl IgG and GBR IgG infusion show intact initial memory encoding: NOR 3 h (Ctrl IgG n=7, GBR IgG n=8) after familiarization (Fam.; ratio of the preference scores: 1.16 ± 0.1 for Ctrl IgG; 1.29 ± 0.04 for GBR IgG; LMM main effect analysis). **c)** Memory consolidation (NOR 24 h; Ctrl IgG n=10, GBR IgG n=9) is impaired in GBR IgG infused mice (ratio of the preference scores: 1.31 ± 0.04 for Ctrl IgG; 1.14 ± 0.06 for GBR IgG; LMM with BH-corrected simple effect analysis). Boxplots show median and interquartile range. *n.s.*, not significant. Statistical details in **Extended Data Table 2**. See also **Extended Data** Figure 2a.

### GABA_B_R IgG specifically interferes with presynaptic GABA_B_R function

To elucidate the impact of anti-GABA_B_R aAbs on synaptic function, and to differentiate their specific effects on GABA_B_Rs located at the pre- and postsynaptic nerve terminals, we performed whole-cell patch-clamp recordings in acute hippocampal slices from mice chronically infused with GBR or Ctrl IgG.

First, we tested the effect on the presynaptic GABA_B_ autoreceptors and recorded pharmacologically isolated, evoked inhibitory postsynaptic currents (eIPSCs) in CA1 pyramidal neurons (**Fig. 3a**). After Ctrl IgG, eIPSC amplitude exhibited a progressive depression during a stimulus train, with the 10^th^ response reduced to 58.7 ± 15.2% of the initial IPSC amplitude. The specific GABA_B_R blocker CGP55845 (10 µM) attenuated this depression, thereby reflecting the GABA_B_R-mediated modulatory effect (**Fig. 3b**, eIPSC_10_/eIPSC_1_ = 70.0 ± 15.9%). These results confirm that presynaptically released GABA can feed back and activate GABA_B_ autoreceptors, thereby enhancing the short-term depression during repetitive stimulation. Conversely, in GBR IgG infused mice, we observed a substantially weaker depression of IPSC amplitudes compared to recordings in artificial CSF (aCSF) (**Fig. 3b**, Ratio IPSC_10_/IPSC_1_ = 67.1 ± 11.6%). This inhibitory effect on GABA_B_Rs was not further altered by application of CGP55845 (eIPSC_10_/eIPSC_1_ = 68.9 ± 10.9%). Thus, GBR IgG affects autoreceptors with an efficiency comparable to chemical inhibition by CGP55845.

**Fig. 3:**
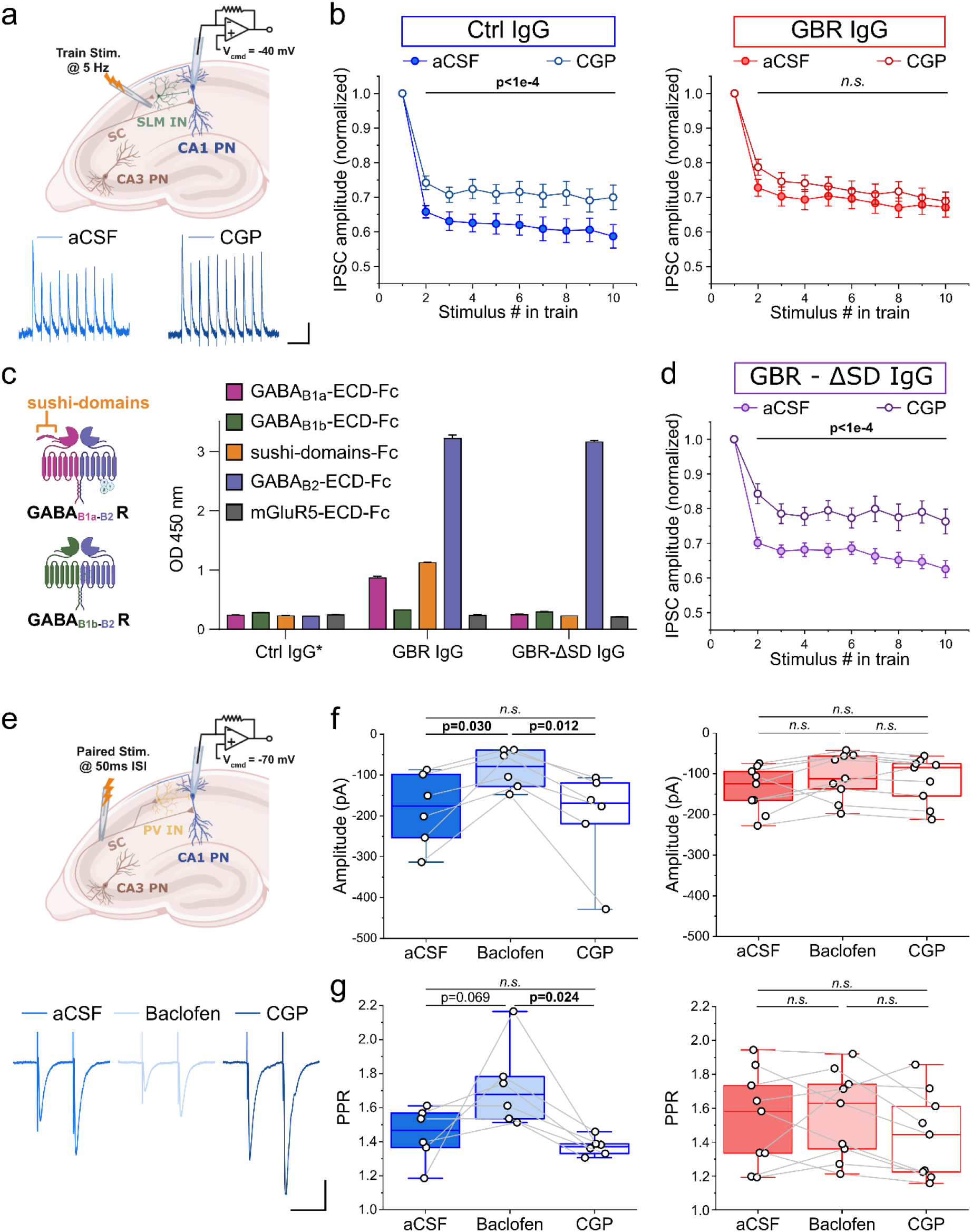
GBR IgG inhibits presynaptic GABA_B_ auto-and heteroreceptors. **a**) Electrophysiological investigation of GABA_B_ autoreceptors. Top: schematic of eIPSC recording in CA1 pyramidal cells (*V_hold_* = −40 mV) evoked by 5 Hz *stratum lacunosum-moleculare* (SLM) stimulation (10 impulses). Bottom: example eIPSC traces before (aCSF; blue) and after (CGP; dark blue) bath perfusion of CGP55845. Scale bars: 500 ms, 20 pA. **b**) eIPSC amplitudes normalized to first eIPSC in train. Left: CGP55845 (CGP) significantly diminishes train-induced eIPSC depression in Ctrl IgG-treated mice (blue circles; n=20). Right: GBR IgG attenuates baseline depression (aCSF; red full circles; n=17), occluding further CGP modulation (red open circles). Two-way repeated-measures [RM] ANOVA with Tukey’s post-hoc tests). **c**) Left: schematic of pre- and postsynaptic GABA_B_Rs containing the GABA_B1a_/SD or the GABA_B1b_ subunit, respectively. Right: Binding of GBR IgG before and after depletion of anti-sushi-domain reactivity (GBR-ΔSD IgG; 50 µg/ml) to soluble Fc-tagged ECDs of the GABA_B_R or of mGluR5 (negative control) as determined by ELISA. Ctrl IgG* was derived from a patient with an unrelated autoimmune disease. GBR-ΔSD IgG lacks reactivity to the GABA_B1a_ subunit and isolated SDs, compared to GBR IgG. **d**) Same as (B), but for GBR-ΔSD IgG-treated mice (violet, n=15). SD-depletion yields Ctrl IgG-like eIPSC depression (aCSF; violet full circles), which is significantly attenuated by CGP (violet open circles). Two-way RM ANOVA. **e**) Electrophysiological investigation of GABA_B_ heteroreceptors. Top: schematic of paired-pulse eEPSC recordings in CA1 pyramidal cells (*V_hold_* = −70 mV) following Schaffer collaterals (SC) stimulation (interstimulus interval [ISI] = 50 ms). Bottom: example traces of eEPSCs in aCSF (blue), in 5 µM baclofen (light blue), and after removing baclofen while washing in 10 µM CGP55845 (dark blue) to the same cell. Scale bars: 50 ms, 50 pA. **f**) First impulse eEPSC amplitudes in Ctrl IgG-infused mice (blue; n=6) get reduced by baclofen, which is reversed by CGP. GBR IgG (red; n=9) prevents this baclofen-mediated eEPSC modulation. **g**) Same as (F), but for short-term plasticity. GBR IgG suppresses the baclofen-induced increase in paired-pulse ratio (PPR: eEPSC_2_ / eEPSC_1_). (f-g) One-way RM ANOVA with Tukey’s post-hoc tests. Curves show mean ± SEM. Boxplots show median and interquartile range. Bars in **C**) show mean ± s.d. *n.s.*, not significant. Statistical details in **Extended Data Table 2**. See also **Extended Data** Figure 1c.

To determine whether the reactivity to the primarily presynaptically localized GABA_B_Rs containing SDs is critical for the effect of GBR IgG on hippocampal synapses in mice, we depleted SD-reactive aAbs from GBR IgG using a recombinant SD protein. The resulting IgG fraction lacked the reactivity to the SDs and to the GABA_B1a_ subunit, whereas the reactivity to the GABA_B2_-ECD and postsynaptic terminals was preserved (**Fig. 3c; Extended Data Fig. 1c**). Interestingly, in acute slices from mice infused with GBR-ΔSD IgG, the function of GABA_B_ autoreceptors on IPSC amplitudes was not impaired: similarly to Ctrl IgG-treated mice, CGP55845 markedly attenuated synaptic depression (eIPSC_10_/eIPSC_1_ aCSF = 62.6 ± 9.6% vs eIPSC_10_/eIPSC_1_ CGP = 76.3 ± 13.8%; **Fig. 3d**). These findings confirm that the effects of GBR IgG were specifically induced by GABA_B1a_/SD-reactive aAbs.

Furthermore, we tested presynaptic GABA_B_ heteroreceptor-mediated inhibition of glutamate release at excitatory synapses by recording pharmacologically isolated evoked excitatory postsynaptic currents (eEPSCs) in CA1 pyramidal neurons (**Fig. 3e**). After Ctrl IgG, bath perfusion of the GABA_B_R agonist baclofen (5 µM) led to a substantial reduction of eEPSC amplitudes, as activation of presynaptic GABA_B_ heteroreceptors inhibits vesicle release, which was again reversed by CGP55845 (**Fig. 3f**). In contrast, in slices from GBR IgG infused mice, baclofen and CGP55845 perfusion had a negligible influence on eEPSC amplitudes (**Fig. 3f**).

Besides, also the short-term synaptic plasticity (calculated by the paired-pulse ratio, PPR: eEPSC_2_ / eEPSC_1_) showed a pronounced facilitation in both Ctrl IgG- and GBR IgG-treated mice (PPR ≍ 1.4–1.6; aCSF), indicating a relatively low release probability (**Fig. 3g**). In Ctrl IgG-infused mice, baclofen markedly increased the PPR, which was reversed by CGP55845, but after GBR IgG neither baclofen nor CGP55845 was able to alter the PPR (**Fig. 3g**). Together, these results suggest that anti-GABA_B_R aAbs bind to GABA_B_ heteroreceptors at glutamatergic terminals and block their activation by agonists like baclofen.

### GABA_B_R IgG does not alter postsynaptic GABA_B_ receptor activity and intrinsic neuronal excitability

Having demonstrated that GBR IgG exert an antagonist-like effect on presynaptic GABA_B_ auto- and heteroreceptors, we next tested whether these aAbs also alter postsynaptic GABA_B_R activity and neuronal excitability. First, we evaluated the impact of GBR IgG on passive neuronal properties and found unchanged subthreshold membrane properties of CA1 pyramidal neurons (**Fig. 4a-b**). In order to test the effects on postsynaptic GABA_B_R signaling, we applied a high-frequency extracellular stimulation (HFS) to the *stratum lacunosum-moleculare* (SLM) to induce GABA release and local activation of postsynaptic GABA_B_Rs (**Fig. 4c**). Since postsynaptic signaling via GABA_B_Rs is driven by GIRK-channel activation^33,34^, we first analyzed the current-voltage (I-V) relationship of slow inhibitory postsynaptic currents (sIPSC) at different holding potentials. This yielded an average reversal potential of −102.7 ± 3.6 mV for Ctrl IgG and −109.1 ± 4.3 mV for GBR IgG treatment groups. These values are close to the calculated potassium equilibrium potential (V_eq_[K^+^] = −104.5 mV) for our solutions, confirming that sIPSCs are induced by K^+^ channels^1^ (**Fig. 4d**). The sIPSPs were proven to be GABA_B_R-mediated by the complete blockade upon CGP55845 and remained unchanged in slices of Ctrl IgG and GBR IgG-infused animals (**Fig. 4e**).

**Fig. 4:**
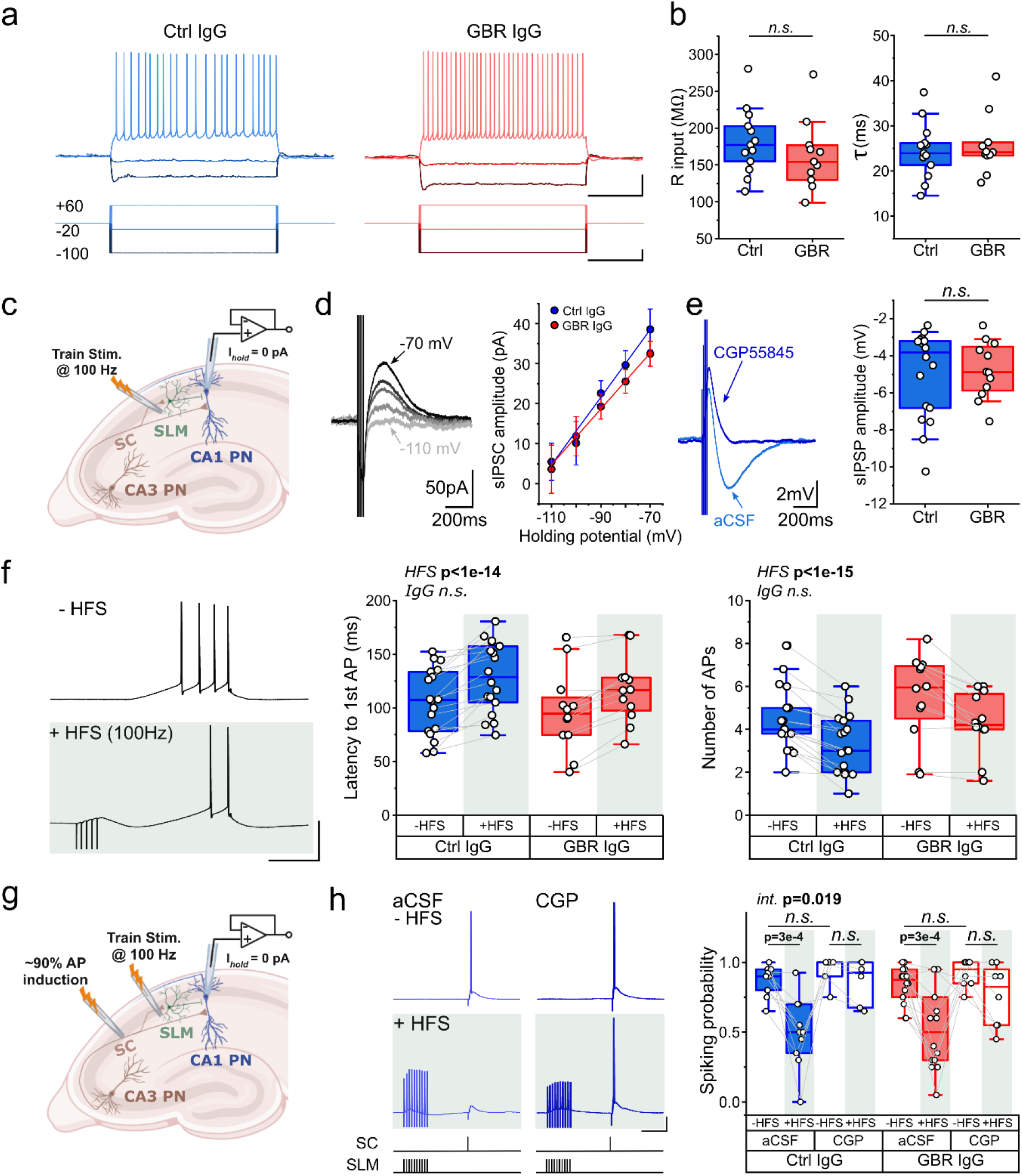
Postsynaptic GABA_B_R influence on GIRK channels and intrinsic excitability remain unaffected. **a**) Example current-clamp recordings of CA1 PCs from Ctrl IgG- (blue) and GBR IgG-treated (red) mice. Bottom: injected current steps (scale bars: 1 s, 50 pA). Top: voltage responses (scale bars: 1 s, 20 mV). **b**) Subthreshold membrane properties. Input resistance (left) and membrane time constant *τ* (right) remain unchanged (Ctrl IgG n=13; GBR IgG n=11; two-sample *t*-test or Mann-Whitney U-test). **c**) Schematic of postsynaptic GABA_B_R recordings in CA1 pyramidal neurons (*I_hold_* = 0pA) triggered by 5-impulses train stimulations at 100 Hz in the SLM. **d**) Current-voltage (I-V) relationship to test the potassium equilibrium potential (V_eq_[K^+^]), confirming the GABA_B_R-dependent GIRK channel activation. Left: Example sIPSCs at holding potentials from −110 to −70 mV (scale bars: 200 ms, 50 pA). Right: I-V reveal a reversal potential close to the V_eq_[K^+^] for both groups (Ctrl IgG: −102.7 ± 3.6 mV, n=13; GBR IgG: −109.1 ± 4.3 mV, n=9).**e**) Left: example slow inhibitory postsynaptic potential (sIPSP) driven by GABA_B_R-mediated GIRK channel activation (aCSF; blue), completely blocked by 10 µM CGP55845 (dark blue; scale: 200 ms, 2 mV). Right: GBR IgG does not affect sIPSP amplitudes (Ctrl IgG n=18; GBR IgG n=12; Mann-Whitney U-test). Recordings were isolated for GABA_B_R-mediated potentials (10 µM CNQX, 50 µM AP-V and 20 µM Bicuculline). **f**) Left: Example AP-inducing current ramp protocol (0–300 pA over 200 ms) without (-HFS) and with (+HFS) a preceding 100 Hz stimulation (scale bars: 100 ms, 50 mV). Middle/right: GABA_B_R activation by HFS increases the latency to 1^st^ AP and reduces the number of APs without differences between Ctrl IgG (n=18) and GBR IgG (n=12). LMM main effect analysis. **g**) Schematic of AP recordings in CA1 pyramidal neurons (*I_hold_* = 0pA) induced by SC stimulation and AP blockade by a preceding 10-impulses train stimulation at 100 Hz in the SLM. **h**) Left: Stimulation protocol and example SC-induced AP-traces (-HFS) following a 250 ms preceding SLM stimulation (+HFS) for a CA1 neuron recorded in naïve aCSF versus 10 µM CGP55845 (scale bars: 100 ms, 20 mV). Right: HFS significantly reduces spiking probability in aCSF; this suppression is prevented by CGP equivalently in both Ctrl IgG (n=10) and GBR IgG (n=14) groups. Grey lines depict paired recordings. Three-level LMM with BH-corrected simple effect analysis. Curves show mean ± SEM. Boxplots show median and interquartile range. *n.s.*, not significant. Statistical details in **Extended Data Table 2**.

To assess effects of anti-GABA_B_R aAbs on neuronal excitability, we utilized two established approaches to study GABA_B_R-mediated changes in excitability. First, we applied a depolarizing ramp protocol to initiate action potentials (AP) in the presence or absence of preceding HFS in SLM for local GABA_B_R activation. HFS delayed the latency to the first AP and reduced the number of APs (**Fig. 4f**) indicating lowered intrinsic excitability and increased threshold of AP initiation due to postsynaptic GABA_B_R-mediated sIPSPs. These findings were equally observed in slices of both, Ctrl IgG- and GBR IgG-treated mice (**Fig. 4f**). Second, to elucidate whether the GABA_B_Rs are still able to regulate the hippocampal output after GBR IgG treatment, we investigated the effect of postsynaptic GABA_B_R-mediated inhibition on the activation of CA1 pyramidal cells. For this, we applied a short impulse to the SC in a stimulation strength sufficient to induce APs in ∼90-95% of the repetitions (**Fig. 4g-h**). A HFS in SLM 250 ms prior to the SC impulse drastically reduced AP firing, which is known as GABA_B_R-mediated spike-blocking efficacy^35^. AP spiking probability was reduced by HFS from 0.86 ± 0.03 (Ctrl IgG, aCSF, -HFS) to 0.50 ± 0.08 (Ctrl IgG, aCSF, +HFS) and reversed by CGP (spiking probability 0.94 ± 0.04 vs 0.86 ± 0.06 for -HFS and +HFS, respectively). Again, spike-blocking efficacy mediated by postsynaptic GABA_B_Rs was preserved after GBR IgG treatment (**Fig. 4h**). Thus, consistent with the unchanged postsynaptic GIRK-mediated hyperpolarization (sIPSP), the physiological modulation of intrinsic excitability, firing threshold, and spike blocking efficacy in CA1 neurons is not altered by GBR IgGs.

### GABA_B_R IgG disinhibits hippocampal pathways and impairs long-term potentiation

We next evaluated GBR IgG effects on hippocampal pathways and long-term synaptic plasticity. Mossy fiber (MF) synapses provide major excitatory input to the CA3 network, where GABA_B_Rs effectively shape these input patterns via heterosynaptic depression to maintain the sparse coding of dentate gyrus (DG) signals^36^. To investigate how GBR IgG influences this MF→CA3 pathway, we recorded field excitatory postsynaptic potentials (fEPSPs) in CA3 *stratum lucidum* upon paired (ISI 40 ms) or single (0.05 Hz) MF stimulation (**Fig. 5a-b**). The paired-pulse ratio (PPR) showed a pronounced facilitation (PPR: Ctrl IgG = 3.56 ± 0.22) verifying that CA3 activation is predominantly driven by MF input. This ratio was reduced in GBR IgG group (2.93 ± 0.14), suggesting an increased release probability onto CA3 pyramidal neurons (**Fig. 5c**). The input-output (I/O) curve of fEPSP amplitudes plotted against the fiber volley (FV) amplitudes revealed a larger postsynaptic output for the same presynaptic input in GBR IgG-treated mice (**Fig. 5d**). Moreover, baclofen prominently reduced fEPSP amplitudes in the Ctrl IgG group, whereas GBR IgG attenuated this baclofen-induced suppression (**Extended Data Fig. 2b**). Next, we assessed fEPSPs in the CA1 *stratum radiatum* (SR) after SC stimulation to evaluate signal transduction from CA3 to CA1 (**Fig. 5e**). Consistent with our findings in the MF-CA3 pathway, I/O stimulation from 0 to 1.0 mA in 0.1 mA steps revealed an increased fEPSP slope in slices from GBR IgG-infused mice (**Fig. 5f**).

**Fig. 5:**
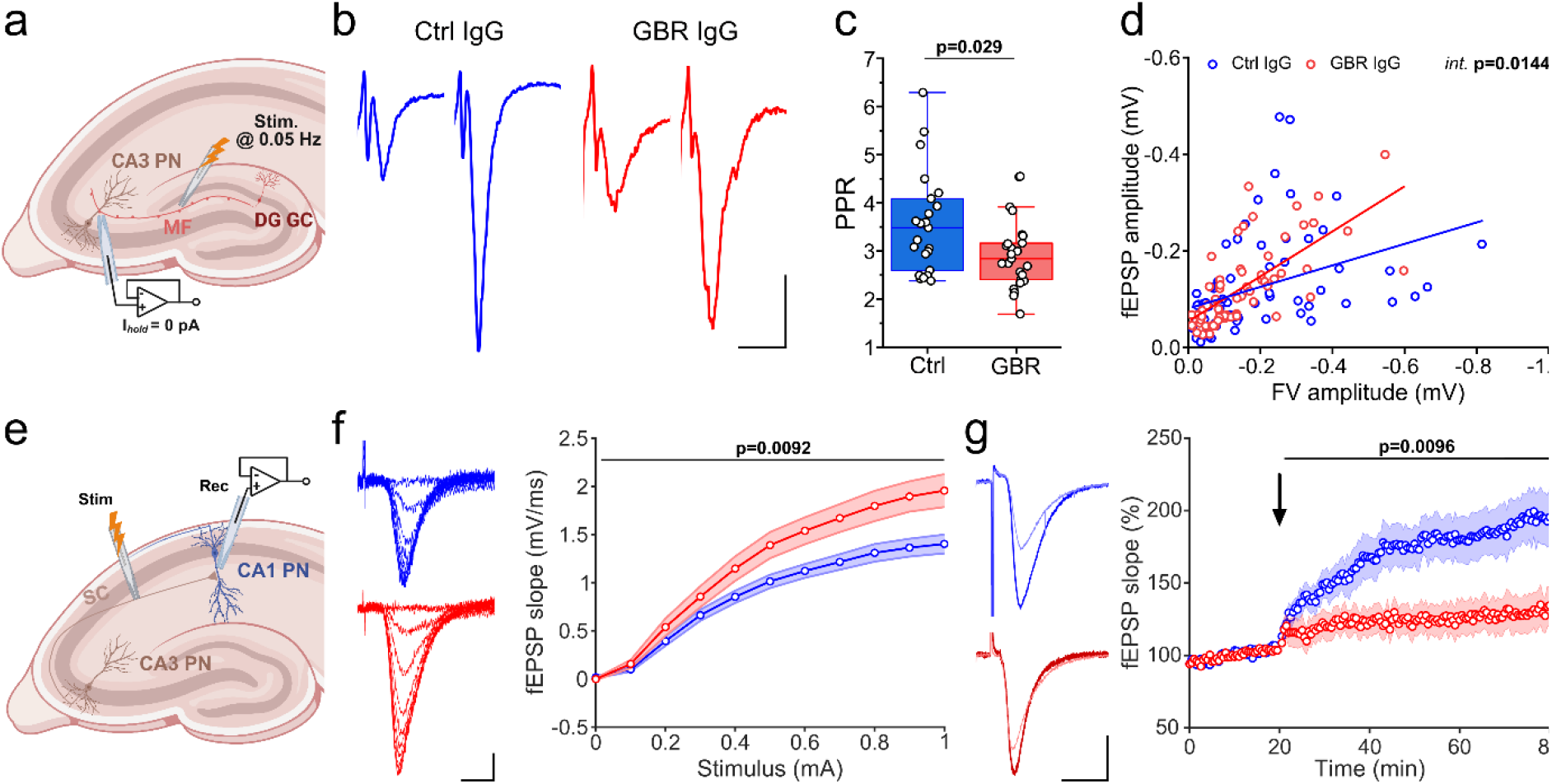
GBR IgG unconstrains network excitability and reduces the induction of synaptic plasticity. **a)** Schematic of mossy fiber (MF) pathway stimulation (0.05 Hz) in the dentate gyrus (DG), and field excitatory postsynaptic potential (fEPSP) recordings in CA3 *stratum lucidum* in *ex vivo* brain slices. **b)** Example paired-pulse recordings (ISI = 40 ms) of Ctrl IgG-(blue) and GBR IgG-treated (red) mice. ISI and stimulation artifact are removed for clarity. Scale bars: 10 ms, 0.2 mV. **c)** Paired-pulse ratio (PPR) showing strong facilitation under Ctrl IgG (3.56 ± 0.22; n=23), which is reduced by GBR IgG treatment (2.93 ± 0.14; n=25; Mann-Whitney U-test) **d)** CA3 Input/Output (I/O) scatter plot of fEPSP versus presynaptic fiber volley (FV) amplitudes (Ctrl IgG n=23; GBR IgG n=21 slices) with linear regression fits. GBR IgG increases postsynaptic output (steeper slope) at constant presynaptic input. Linear regression with interaction term; slope difference *p*=0.014, intercepts difference *p*=0.21. **e)** Schematic of SC pathway stimulation and fEPSP recordings in CA1 *stratum radiatum* (SR). **f)** Left: Example SC-induced fEPSPs at different stimulation steps in Ctrl IgG (blue) and GBR IgG (red) slices (scale bars: 5 ms, 0.5 mV). Right: CA1 I/O curves (0–1.0 mA, in 0.1 mA steps) show increased network excitability in GBR IgG-treated mice (whole-curve permutation test; Ctrl IgG n=31, GBR IgG n=32 slices). **g)** Left: Example fEPSPs before (thin) and after TBS (bold; scale bars: 5 ms, 0.5 mV). Right: CA1 fEPSP time courses demonstrating robust increase of fEPSP slopes after TBS induced LTP in Ctrl IgG-treated mice compared to GBR IgG (whole-curve permutation test; Ctrl IgG n=11, GBR IgG n=17). Arrow indicates the onset of LTP induction. Curves show mean ± SEM. Boxplots show median and interquartile range. Statistical details in **Extended Data Table 2**. See also **Extended Data** Figure 2b-c.

GABA_B1a_R knockout mice show defective memory consolidation^32^, comparable to the results we obtained upon intraventricular passive transfer of GBR IgG (**Fig. 2**). Moreover, both GABA_B1a_R knockout^11^ and acute pharmacological GABA_B_R antagonism^37,38^ impair hippocampal long-term potentiation (LTP). Therefore, we next tested GBR IgG effects on synaptic LTP in the CA3→CA1 pathway. LTP was induced using theta-burst stimulation (TBS) of SC, where CA1 fEPSPs were recorded for 60 minutes post-TBS and revealed a pronounced reduction in LTP strength in GBR IgG-treated mice (**Fig. 5g**). In a separate set of recordings, we found that this LTP reduction was not further affected by CGP55845 (10 µM), thereby corroborating the antagonist-like effect of aAbs on GABA_B_Rs (**Extended Data Fig. 2c**).

Together, these results indicate that ambient GABA mediates a tonic heterosynaptic depression in these pathways, which is disrupted by the blocking effects of anti-GABA_B_R aAbs on GABA_B_R function. This leads to defective synaptic plasticity in the SC→CA1 pathway and recapitulates the synaptic and plasticity pathophysiology observed in models of GABA_B1a_R deficiency.

### GBR IgG induces hippocampal hyper-coupling and amplifies CA1 network oscillations *in vivo*

As GABA_B_Rs modulate fast network activity and rhythmogenesis through their slow dynamics, our data showing defects in GABA_B_R-mediated synaptic transmission and plasticity *ex vivo* led us to analyze hippocampal network oscillations and coupling *in vivo* in freely behaving mice—processes critical for memory processing and consolidation^39–44^. We recorded simultaneously from CA1 and CA3 areas *in vivo* after intra-hippocampal injection of either Ctrl IgG or GBR IgG (**Fig. 6a** and see methods for details). The local field potential (LFP) signals *in vivo* were then quantified for network oscillations and the interareal coupling between the phase of CA3 theta oscillations and the amplitude of CA1 gamma oscillations, using the modulation index^45,46^. This measure quantifies the degree to which gamma amplitude preferentially occurs at specific phases of the theta cycle (MI; **Fig. 6a**). Under Ctrl IgG, the coupling was mainly confined to the CA3 phase in 6–10 Hz theta band and the CA1 amplitude in 50–85 Hz high-gamma band (**Fig. 6b-c, Extended Data Fig. 3a**), consistent with previous reports^40,47–49^. GBR IgG abnormally increased the coupling strength (peak MI; **Fig. 6c** [left]), without affecting the core theta and gamma frequencies of coupling (**Extended Data Fig. 3a**). However, this hyper-coupling was more spectrally focused, as GBR IgG reduced the overall coupling frequency-area (**Fig. 6c** [middle]), through narrowing of both the theta pacemaker bandwidth and the gamma carrier bandwidths (**Fig. 6c** [right], **Extended Data Fig. 3b**). Together, GBR IgG pathologically locks the hippocampal circuit into a less flexible, spectrally rigid state of CA3→CA1 communication.

**Fig 6.**
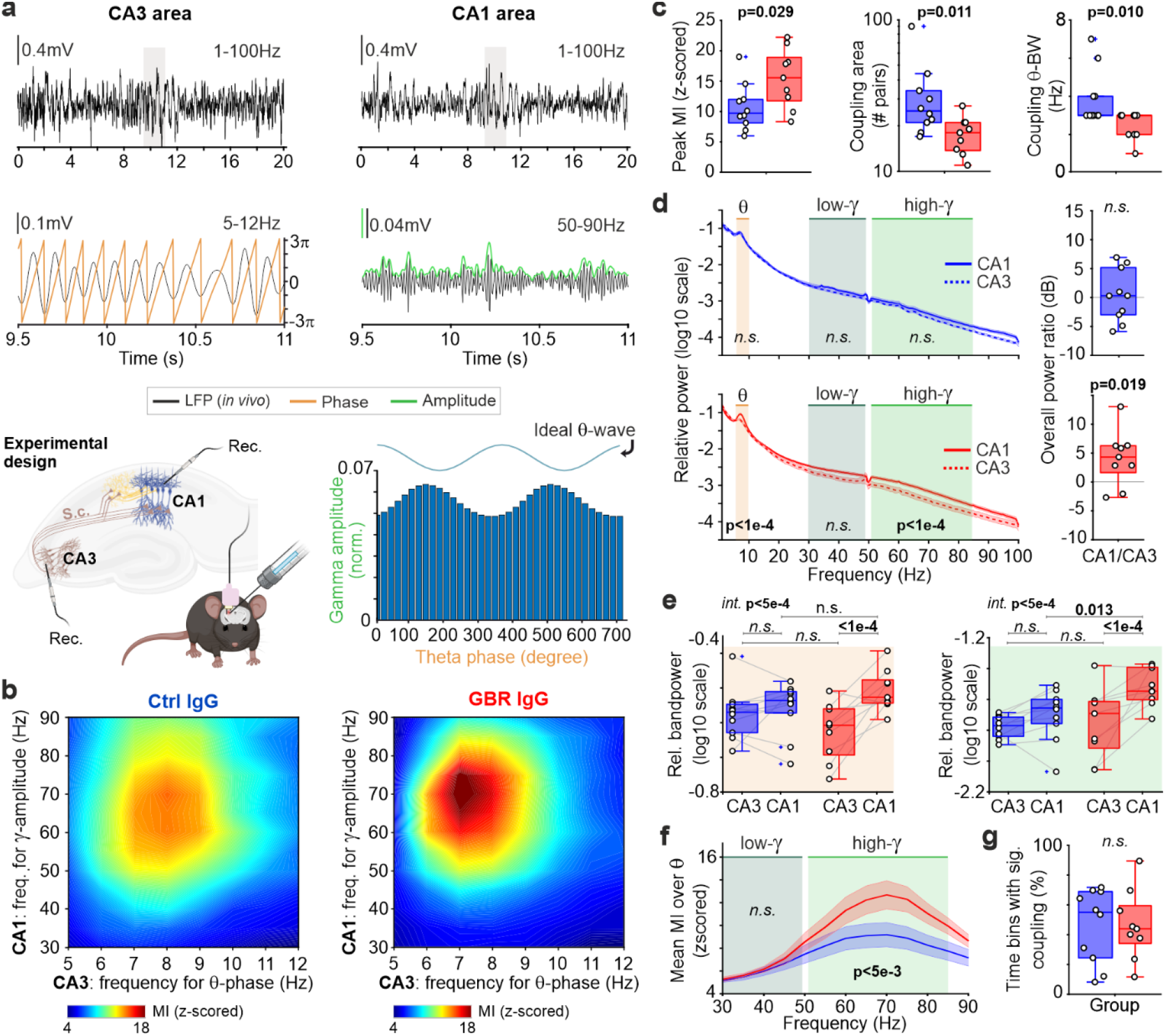
GBR IgG induces hippocampal hyper-coupling and amplifies CA1 network oscillations *in vivo*. **a)** Top: example broad-band LFP recordings. Middle: extracted CA3 theta phase (left) and CA1 gamma amplitude (right). Bottom left: schematic of simultaneous CA3 and CA1 LFP recordings in freely moving mice following intra-hippocampal injection of Ctrl IgG (n=9 mice) or GBR IgG (n=10). Bottom right: resulting phase-amplitude distribution used for modulation index (MI) calculation. **b)** Representative MI comodulograms for Ctrl IgG (left) and GBR IgG (right). The color scale represents z-scored MI, showing stronger coupling in example GBR IgG mouse. **c)** A more spectrally rigid coupling state under GBR IgG. GBR IgG (red) increases peak z-scored MI (left), reduces frequency area of coupling hotspot (middle), mainly via narrowing theta coupling-bandwidth (right) compared to Ctrl IgG (blue) (two-sample *t*-test [left]; Mann-Whitney U-test [middle and right]). **d)** Left: mean CA1 (solid) and CA3 (dashed) relative power spectra in Ctrl (top, blue) and GBR IgG (bottom, red) groups. Shaded areas indicate analyzed frequency bands. Right: median CA1/CA3 spectral ratio, quantifying inter-regional power imbalance (one-sample *t*-test [top]; Wilcoxon signed-rank test [bottom]). The p-values for theta and high-gamma bands stem from (E), and for low-gamma band from **Extended Data** Fig. 3c. **e)** Relative bandpowers in theta (6–10 Hz; left) and high-gamma (50–85 Hz; right) bands. Generalized LMM with BH-corrected simple effects. **f)** Mean z-scored MI averaged over theta (6–10 Hz) plotted versus gamma amplitude frequency. GBR IgG amplifies coupling in the high-but not low-gamma band (Generalized LMM with BH-corrected simple effects). **g)** Percentage of 1-minute time bins (30-s step) per mouse exhibiting significant coupling (MI > 99^th^ percentile versus bin-specific circularly shifted surrogates (two-sample *t*-test). Curves show mean ± SEM. Boxplots show median and interquartile range. *n.s.*, not significant. Statistical details in **Extended Data Table 2**. See also **Extended Data** Figure 3 **and Extended Data** Figure 4.

To investigate the underlying circuit mechanism of this phenomenon, we analyzed the spectral profile of CA1 and CA3 LFPs. This analysis revealed a profound and frequency-specific alteration of the hippocampal circuit’s oscillatory dynamics in the GBR IgG group (**Fig. 6d-e, Extended Data Fig. 3c-d**), as follows: GBR IgG altered the power relationship between the two areas in both the theta (6–10 Hz) and high-gamma (50–85 Hz) bands (**Fig. 6d**). Under Ctrl IgG, the power in these key coupling bands was comparable between CA1 and CA3 (**Fig. 6d**, first row). In contrast, GBR IgG elevated CA1 power relative to CA3 in both frequency bands (**Fig. 6d**, second row), demonstrating a pathological breakdown of the physiological relationship between the source and receiver regions. Notably, CA1 high-gamma power under GBR IgG was the strongest across all tested regions and treatments (**Fig. 6e, Extended Data Fig. 3d**). This spectral specificity was further highlighted by observing unaffected low-gamma band powers (30-50 Hz; **Extended Data Fig. 3c**). Crucially, this frequency-dependent dissociation in power was mirrored by the coupling dynamics: the aberrant hyper-coupling was strictly confined to the high-gamma band, while modulation of the low-gamma band remained unchanged (**Fig. 6f**).

Despite the comparable CA1 theta power across groups (**Fig. 6e** [left]), GBR IgG pathologically amplified CA1 theta power relative to CA3. We considered whether this regional disparity might stem from the difference in intrinsic, sub-threshold resonance properties^50,51^. Indeed, consistent with established biophysical properties^50–53^, CA1 pyramidal neurons from Ctrl IgG group displayed a clear resonance peak in the theta band *ex vivo*, whereas CA3 neurons functioned as low-pass filters (**Extended Data Fig. 4a-c**). Crucially, these characteristic impedance profiles remained unaltered by GBR IgG in both regions (**Extended Data Fig. 4d-e**). Therefore, considering the aAb-induced excessive, upstream excitatory drive (at both MF→CA3 and SC→CA1 synapses; **Fig. 5**), our data suggest that, unlike in CA3, where low-pass filtering tends to dampen the increased drive, the intrinsic theta resonance of CA1 neurons can abnormally amplify this excessive synaptic drive. Consequently, CA1 neurons may pathologically be more effectively entrained by theta rhythms, thereby driving the network-level power imbalance in the theta band.

Finally, we asked whether an increased incidence of coupling events could also contribute to the observed hyper-coupling. However, quantifying the fraction of 1-minute time bins with a significant MI (as compared to surrogate data) showed no difference between groups (**Fig. 6g**). Therefore, GBR IgG does not change the overall temporal incidence of coupling events, but rather profoundly alters the physiological characteristics of the coupling state.

In sum, these results indicate that GBR IgG spectrally confines and strengthens CA3-CA1 coupling by amplifying CA1 network oscillations, which establishes its pathological dominance over the source CA3 area in both the theta and high-gamma bands.

### Network modeling suggests that amplified gamma results from the synergy between hyperexcitability and augmented feedback inhibition

Finally, we sought to investigate the mechanistic link between the altered synaptic kinetics (**Figs. 3, 5**) and the aberrant CA1-network gamma oscillations (**Fig. 6d**). We incorporated our measurements into a biophysical CA1 model comprising pyramidal cells (E), fast-spiking parvalbumin-positive interneurons (I-cells; PV^+^), and O-LM interneurons (O-cells; SOM^+^)^54–56^. Functionally, the E-I subnetwork primarily generates gamma oscillations, while O-cells modulate the network in the theta band. This model (Control) emulates a canonical CA1 microcircuit generating representative theta-nested gamma oscillations (**Fig. 7a**), thus providing a robust framework to interrogate the underlying network dynamics^53,56–58^. To model the GBR IgG effects (GABA_B_R-AB model), we increased both the local inhibitory synaptic weights onto E-cells (*G_OE_* and *G_IE_*) and the extrinsic excitatory drive to E-cells (E-drive; *ε_E_*) by 30%, accounting for our observed increase in the IPSCs and fEPSPs slope (**Figs. 3b and 5f**). Strikingly, this model exhibited a marked amplification in gamma power with preserved theta oscillations (**Fig. 7b-c**), corroborating our *in vivo* findings (**Fig. 6d-e**).

**Fig 7.**
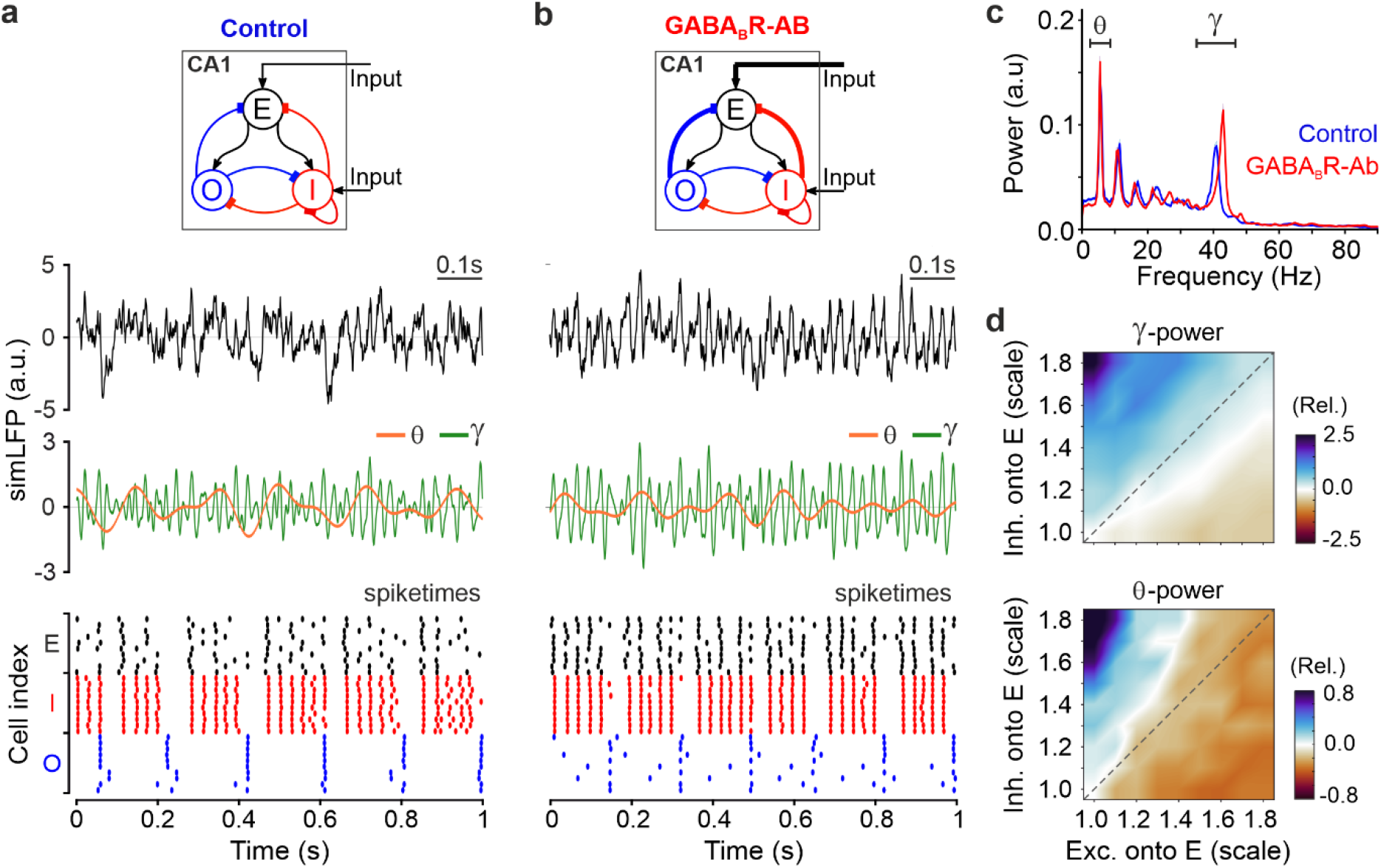
Network modeling suggests that amplified gamma results from the synergy between hyperexcitability and augmented feedback inhibition. **a, b)** Schematic of CA1 network model under Ctrl IgG (a) and GBR IgG (b) conditions (Control and GABA_B_R-AB model, respectively). In the GABA_B_R-AB model, excitatory drive (*ε_E_*) and inhibitory synaptic weights (*G_OE_* and *G_IE_*) onto E-cells were both 30% higher (thick arrows) to mimic hetero- and auto-GABA_B_R dysfunction, respectively. Middle panels show simulated local field potentials (simLFP, black) filtered in theta (orange; 4-10 Hz) and gamma (green; 30-90 Hz) bands. Bottom panels show spike raster plots for pyramidal (E, black), fast-spiking PV^+^ (I, red), and O-LM (O, blue; SOM^+^) cells. Note the tight vertical co-alignment of E- and I-cell spikes into hypersynchronous stripes in (b). **c**) Power spectra of simLFPs. The GABA_B_R-AB model (red) exhibits a marked amplification of representative gamma oscillations (∼42 Hz) compared to Control (blue), while theta power remains comparable. Curves represent mean ± SEM. N=50 trials (trial duration = 10 s). **d**) Parameter-space analysis of network oscillatory dynamics. Heatmaps show the relative change in peak gamma power (top) and theta power (bottom) as a function of excitatory drive (*ε_E_*; X-axis) and synaptic inhibition (both *G_OE_* and *G_IE_*; Y-axis). All values represent the relative change compared to the Control model (coordinate 1.0, 1.0). See also **Extended Data** Figure 5.

To decipher the pathomechanism on the cellular level we analyzed the underlying neuronal spiking. The GABA_B_R-AB model exhibited a hyperexcitable state, with E-cells firing increasing markedly from 15.5 Hz to 19.6 Hz on average (**Fig. 7a-b**, compare black rasters). This activity was pathologically more organized: E-cells showed notably higher spike synchrony (STTC^59^ for [E-E]: 0.49 vs 0.33 in Control; **Extended Data Fig. 5a**). Furthermore, the firing correlation between E- and I-cells, the primary generators of the gamma rhythm, was markedly increased (STTC for [E-I]: 0.51 vs 0.43 in Control **Extended Data Fig. 5a**). These results suggest that the increased feedback inhibition enforces a tighter E-I synchrony, which structures the excessive excitation into coherent, amplified gamma rhythms.

To determine the boundaries of this state, we systematically varied the E-drive and synaptic inhibition from baseline (1.0x) to pathological levels (1.8x) demonstrating that gamma amplification is strictly confined to the upper-diagonal regime where synaptic inhibition is high (**Fig. 7d**). Increasing E-drive alone actually reduced gamma power (**Fig. 7d** [top], lower-diagonal; **Extended Data Fig. 5b-c**) despite increasing firing rates (**Extended Data Fig. 5e**). This indicates that without concomitant increase in inhibition strength, the feedback loop fails to effectively entrain the excessive excitation (**Extended Data Fig. 5f**, right). Consequently, while E-cells may fire in closer proximity to one another, their activity is only loosely structured by interneuronal feedback (**Extended Data Fig. 5b**; bottom), resulting in a reduced population-level gamma power (**Extended Data Fig. 5c**, yellow). Conversely, increasing inhibition alone created a low-firing, slow-frequency state (**Extended Data Fig. 5c-d, g**). Thus, the modeling suggests that the GABA_B_R-AB phenotype requires a “push-pull” synergy: increased extrinsic E-drive prevents the network from being silenced by inhibition, while the increased local inhibition provides the necessary temporal gating to entrain that excessive excitation into a hypersynchronous rhythm.

## Discussion

We leveraged specific pathogenic aAbs against GABA_B_Rs occurring in autoimmune encephalitis to uncover the mechanistic cascade from impaired GABA_B_Rs to dysfunction of synaptic and neuronal signaling, hippocampal circuit dynamics *in vivo*, and ultimately, defective memory consolidation.

### Epitope specificity and mode of action

While previous studies have mapped the binding of polyclonal anti-GABA_B_R aAbs to the N-terminal region of GABA_B1_ subunits^12^, we identified a dual reactivity pattern of patient-derived polyclonal anti-GABA_B_R aAbs to the GABA_B1a_ and GABA_B2_ subunits. This binding pattern is functionally important because the SDs interact with axonal transport proteins and determine the localization of GABA_B_Rs to presynaptic terminals while GABA_B1b_-containing GABA_B_Rs are located at postsynaptic sites^11,19,60^.

Unlike the typical crosslinking and subsequent internalization observed for aAbs against NMDA-, AMPA-, and glycine receptors^27,61,62^, GABA_B_R surface expression remained unchanged, consistent with previous reports^29^. Instead, patient aAbs exerted direct functional antagonism, inhibiting GABA_B_R activation by ambient GABA or the exogeneous agonist, baclofen. This suggests either direct aAb covering of the agonist binding site^63^, or steric hindrance of receptor signaling structures^28^ resulting in interference with agonist-induced receptor signaling. Importantly, when we depleted GABA_B1a_-reactive IgG from the polyclonal fraction, the remaining GABA_B2_-specific aAbs did not alter GABA_B_R function. This reinforces that the observed GABA_B_R dysfunction arises from GABA_B1_ blockade rather than from binding to immunogenic extracellular regions of GABA_B2_, leaving intracellular G-protein signaling via GABA_B2_ preserved. Interestingly, prominent GABA_B2_ reactivity was only observed in patients with paraneoplastic disease, indicating tumor-induced molecular mimicry effects resulting in an additional functionally silent IgG fraction.

### Presynaptic pathology of anti-GABA_B_R aAbs and consequences for cellular signaling and long-term plasticity

GABA_B_Rs regulate presynaptic transmitter release and postsynaptic excitability^20,64^. In line with our biochemical data, our patient IgG induced predominantly presynaptic pathology. Autoantibodies blocked presynaptic auto- and heteroreceptors, abnormally increasing transmitter release and fEPSP amplitudes in CA3 and CA1 areas, phenocopying GABA_B1a_ deficiency^11,25,65,66^. This presynaptic blockade disrupts the heterosynaptic depression important for constraining excitatory transmission and controlling hippocampal sparse coding that regulates mnemonic processes^36,66,67^.

Furthermore, patient IgG markedly reduced hippocampal LTP *ex vivo*. Physiologically, GABA_B_ autoreceptors at interneuron-to-pyramidal terminals reduce GABA release and facilitate NMDAR-dependent LTP^37,68^. Concurrently, GABA_B_ heteroreceptors regulate glutamate vesicles release, maintaining plasticity within a physiological dynamic range and preventing LTP saturation^11^. Accordingly, the absence of these heteroreceptors has been shown to reduce both LTP and the number of silent synapses^11,69^. Therefore, the observed aAb-mediated blockade of GABA_B_ auto- and heteroreceptors may restrain these permissive conditions, directly impairing LTP.

### Hippocampal hyper-coupling and amplified CA1 gamma oscillations as a consequence of autoimmune GABA_B_R dysfunction

Through their slow dynamics, GABA_B_Rs modulate fast network activity, excitatory-inhibitory balance, rhythmogenesis, and information flow in cortical and hippocampal networks ^20,42,44,70–72^. Consistently, our *in vivo* data demonstrated that GABA_B_R aAbs pathologically amplify local CA1 rhythms, thereby disrupting the CA3–CA1 communication. Because the intrinsic excitability and subthreshold resonance of pyramidal neurons remained unaffected, this rhythmopathy is mainly driven by synaptic and network-level dysfunction. Corroborating this, our data-informed computational model identified a specific, intuitive pathomechanism: amplified CA1 gamma power arises from the synergy between unconstrained excitatory drive (driven by impaired heteroreceptors) and augmented feedback inhibition (driven by impaired autoreceptors). Moreover, because CA1, but not CA3, pyramidal neurons intrinsically resonate at theta frequencies (see also^50–53^), they are likely to disproportionately amplify this unconstrained upstream drive at theta frequencies. Ultimately, as our data showed, the CA1 network may functionally escape physiological modulation and lock into a spectrally rigid, hyper-coupled state with CA3, thus reducing the bandwidth for dynamic information transfer.

### Pathological consequences of presynaptic GABA_B_R dysfunction for long-term memory

The observed presynaptic GABA_B_R dysfunction likely underlies the cellular correlates of memory impairment and seizure susceptibility symptoms in anti-GABA_B_R AE^12,13^. Consistent with limbic dysfunction, chronic passive transfer of patient aAbs selectively impaired hippocampal-dependent long-term memory maintenance in our mouse model. This behavioral phenotype closely aligns with previous GABA_B1a_ knockout mouse models^11,32,73^, which exhibited notable defects in long-term object recognition despite largely unaffected initial memory acquisition^32^. Moreover, the reduced hippocampal LTP we observed *ex vivo* likely contributes directly to this diminished precision of long-term memory, as the dynamic range required for effective LTP is compromised.

Physiologically, memory consolidation and retrieval require highly flexible, transient theta-gamma coupling to dynamically coordinate precise neural computations^39–41,43,49^. Therefore, the rigid hippocampal hyper-coupling and amplified CA1 oscillations likely degrade sparse coding capacity and restrict the rich repertoire of hippocampal activity patterns. Consequently, the disrupted theta-gamma coupling may explain specific vulnerability of memory maintenance: while a hyper-coupled state may permit the initial encoding, it impairs the computational flexibility required to stabilize hippocampal engrams and retrieve distinct memory traces over time^74–76^. Indeed, similar alterations in network oscillations are prominent pathomechanisms in other neurological and psychiatric disorders like Alzheimer’s disease^77–79^. Thus, the aAb-mediated GABA_B_R dysfunction presumably acts as a critical switch, stripping the hippocampus of its computational flexibility and driving the profound memory retention deficits of anti-GABA_B_R AE.

### Limitations of the study

First, our use of *in vivo* passive-transfer models was specifically designed to isolate the direct pathophysiological consequences of patient aAb binding on synaptic and network function, independent of systemic inflammation. Therefore, cytotoxic T cell-mediated effects that might contribute to GABA_B_R encephalitis are not modeled here. Second, while our assays conclusively localize the immunogenic regions to the GABA_B1_ SDs and GABA_B2_ extracellular domain, the exact binding epitopes driving its functional antagonism remain to be resolved by high-resolution structural characterization, e.g. by cryo-EM with monoclonal aAbs, as shown for GABA_A_R and NMDAR IgGs^28, 63^. In our experiments using purified patient IgG targeting GABA_B_Rs we capture the comprehensive, clinically relevant pathophysiological complexity of the disease and observed a specific presynaptic pathology. Nevertheless, our findings do not preclude additional pathogenic effects arising from other aAbs, not detected in our assays, that target distinct epitopes of GABA_B1_ or the fully assembled GABA_B_R. Lastly, since we here focused on synaptic and perisynaptic GABA_B_Rs, the effects of IgG on extrasynaptic dendritic GABA_B_Rs and on GABA_B_Rs on glial cells remains to be determined.

Overall, our results position GABA_B_R dysfunction as a mechanistic link between autoimmunity, circuit dysregulation, and cognitive decline, thereby establishing immune-mediated synaptic pathology as an alternative mechanistic pathway to dementia disorders, distinct from classical neurodegenerative processes.

## Resource availability

### Lead contact

Further information and requests for resources and reagents should be directed to and will be fulfilled by the lead contact, Christian Geis.

### Materials availability

GBR IgG and GBR-ΔSD IgG was created during this study and is available upon reasonable request. Due to the finite available quantity, the patient material used in this study may be available on a limited basis from the lead contact with a completed Materials Transfer Agreement.

### Data and code availability

Statistical data associated with this study is presented in Extended Data Table 2. This paper does not report original code. HOC code associated with the neural network model is publicly available in ModelDB github (https://github.com/ModelDBRepository/138421). The original Matlab code for phase-amplitude coupling (https://github.com/tortlab/phase-amplitude-coupling) and EEGLAB toolbox (https://sccn.ucsd.edu/eeglab/) used for filtering signals are also available online. Any additional information required to reanalyze the data reported in this work is available from the lead contact upon request.

## Supporting information

Extended Data Table 1 containing patient characteristics.

Extended Data Table 2 containing synopsis of statistical tests.

## Acknowledgements

We thank Claudia Sommer, Christin Reißig, and Cathrin Bayer-Kaufmann for excellent technical assistance. For administrative tasks and the coordination of animal transport between collaborating facilities, we would like to thank Anja Strauß, Melanie Brese, and Anke Schönherr. This work was supported by ‘Zentrale Forschungswerkstätten Jena’ (Central Research Workshops) of Jena University Hospital with production of relevant laboratory and electrophysiology equipments. This work was supported by THiMEDOP.

## Author contributions

J.S., E.A.L. and C.G. conceived the project and designed the experiments. E.A.L. carried out the behavioral experiments. J.S. and M.M. performed all patch-clamp electrophysiology experiments, E.A.L., A.S., T.G., D.S. the *ex vivo* field potential recordings, and E.A.L. together with H.H. the *in vivo* LFP recordings. E.A.L. and T.K. did the imaging experiments, under supervision of S.H. V.R. established an analysis pipeline for *in vivo* LFP recordings, analyzed the network oscillations and created the biophysical network model. C.F., F.L., S.J., N.M., J.L., A.K., and H.P. provided patient samples and clinical data. H.-C.K. tested the subunit reactivity of patient sera using ELISA and depleted SD-reactive antibodies from GBR IgG. H.-C.K., D.S., C.S.-H., H.P., C.F., and C.G. provided valuable input and expertise to the experiments and interpretation. J.S., E.A.L., V.R., and C.G. wrote the manuscript and designed figures. All authors reviewed and edited the manuscript. C.G., H.-C.K., C.S.-H. acquired funding. C.G. supervised all aspects of the study.

## Methods

### Experimental model and subject details Human IgG samples and preparation

Patient-derived samples of serum were acquired after informed consent and local ethics approval. Control IgG samples were acquired from patients with an unrelated autoimmune disease without antineuronal reactivity (Ctrl IgG and Ctrl IgG*). The patient-derived GBR IgG fraction was purified from plasmapheresis filtrates of patients by Protein G Sepharose 4 Fast Flow (Cytiva Europe GmbH) as described previously^80^.

### Primary cell cultures

Primary hippocampal neurons were isolated from E18 embryos of 10 female C57BL/6 mice. The genotype of the embryonic cells was not determined, and the cultures presumably contained both sexes. Animal breeding and housing were performed according to the ARRIVE guidelines for reporting animal research^81^, and all animals were approved by the local ethics committee at the University of Jena (#TWZ-08-2020 and #UKJ-23-002). The preparation of the E18 tissue was performed as described earlier^82^. Briefly, embryonic brains were removed, and hippocampi separated, trypsinated in 0.25% trypsin solution and triturated. Neurons were first plated on poly-D-lysin-(PDL) coated coverslips in plating minimum essential medium (MEM, 0.5% glucose, 10% horse serum, 0.5% penicillin/streptavidin) with a density of 20.000 cells/cm^2^. After 1 h plating medium was replaced by growth medium (Neurobasal, 2% B27 [Life Technologies], 1% glutamine, 0.5% penicillin/streptavidin) and neurons were cultured for 10−14 days before use.

### Mice

96 C57BL6/J mice, ten to twenty weeks old were used in this study. Animal experiments were performed according to the ARRIVE guidelines for reporting animal research^81^, and the experimental protocol was in accordance with European regulations (Directive 2010/63/EU) and was approved by the local ethics committee at the University of Jena (TVA #UKJ-17-053, #UKJ-22-029 and #UKJ-23-002). All experimental and breeding animals were kept under a 14:10 hours day:night cycle with water and food *ad libitum*. Animal experiments were performed during the light phase of the cycle. Behavioral and electrophysiological experiments were performed after cerebroventricular infusion or direct intrahippocampal injection of human polyclonal IgG fractions with either GBR IgG or non-reactive Ctrl IgG.

## Method details

### ELISA detection of anti-GABA_B_ receptor IgG in patient serum

The reactivity of patient sera or IgG fractions to extracellular domains of the GABA_B_ receptor was tested using an ELISA-based approach^83,84^. Briefly, rabbit Fc fusion proteins of the extracellular domain (ECD) of GABA_B1a_ (GenBank accession no. NM_001470, amino acids 1-590), the sushi-domains (SDs) of GABA_B1a_ (amino acids 1-164), the ECD of GABA_B1b_ (NM_021903, amino acids 1-473), the ECD of GABA_B2_ (NM_005458, amino acids 1-480) and of the ECD of mGluR5 (NM_001143831, amino acids 1-580) were expressed in HEK293T cells and immobilized onto 96-well plates via anti-rabbit IgG, Fc fragment specific (111-005-046, Jackson ImmunoResearch, 10 µg/ml). Sera or IgG fractions were diluted in StartingBlock (PBS) Blocking Buffer (37538, Thermo Fisher Scientific)/0.05% Tween-20 and IgG binding was detected using HRP-conjugated anti-human IgG (709-036-149, Jackson ImmunoResearch) and 1-step Ultra TMB ELISA substrate (34028, Thermo Fisher Scientific).

For the heat map presentation, we calculated net absorbance values at 450 nm based on IgG binding to the GABA_B_R antigens by subtracting the individual background value on mGluR5-ECD for each serum/IgG and normalized them to the value of GBR-1 IgG on GABA_B2_-ECD in the specific assay.

### Depletion of anti-sushi domain reactivity from GBR IgG

The SDs of GABA_B1a_ (amino acids 1-164) fused to a C-terminal Twin-Strep-tag were expressed in HEK293T cells and immobilized on Strep-Tactin XT beads (2-5030-010, iba lifesciences). GBR IgG was passed over a column containing the SD-coated beads, collected, quantified and tested for remaining anti-GABA_B_R reactivities using ELISA.

### Immunostainings and Confocal Laser Scanning Microscopy (LSM)

To evaluate the effective binding of the patient derived IgG or serum with AB to the GABA_B_R, co-stains of the receptor and the human IgG were performed in primary cell cultures. After 14 days *in vitro* (DIV), neuronal cultures were incubated with 10 µg/ml solution of patient derived IgG fractions or patient serum in a 1:50 dilution for 30 minutes at 37 °C, followed by 4% paraformaldehyde (PFA) fixation for 20 minutes at room temperature (RT). Cells were blocked with 10% normal goat serum and 10% bovine albumin serum in phosphate buffered saline (PBS) 1 h at RT, then a 1:200 dilution of secondary antibody goat anti-human Alexa Fluor 488 (Invitrogen) was applied for 2 h at RT. Subsequently, a second round of blocking and permeabilization with 0.1% Triton was performed for 1 h at RT, then cell cultures were incubated overnight at 4 °C with either one of these primary AB against the GABA_B_R: 1) a commercially available polyclonal rabbit AB against the GABA_B2_ subunit 1:200 (Synaptic System); or 2) a recombinant, custom-made polyclonal rabbit AB against the GABA_B1_ subunit 10 µg/ml^26^, kindly provided by Prof. Shigemoto (IST, Austria). Afterwards, cells were incubated with secondary goat-anti-rabbit Alexa Fluor 647 antibodies (Invitrogen) 1:200 for 2 h at RT. Lastly, cells were stained with 4′,6-diamidino-2-phenylindole (DAPI) for 5 minutes, washed with PBS, and mounted on glass slides for the confocal microscope acquisition.

LSM 980 Zeiss confocal microscope (Carl Zeiss Microscopy GmbH, Jena) was used to acquire images of the immunostained neurons using laser lines for excitation of Alexa Fluor 647 and 488 (642 nm and 488 nm, respectively). All recording parameters were kept constant during acquisition (pinhole at 5.0 Airy Unit (AU)/328 µm; laser power 488 nm at 5.0% and 642 nm at 1.0%; detector gain at 850 V; pixel size at 0.043 µm x 0.043 µm x 0.170 µm; 16-bit depth). Images were analyzed and pixel intensity-based colocalization of antibody-derived fluorescent signals was performed and expressed as Pearson’s Coefficient.

### Internalization Assay and Confocal LSM

To assess the effect of aAbs on receptor surface expression, we performed a receptor internalization assay. Neuronal cultures at 14 DIV were incubated for 24 h at 37 °C with 10 µg/ml of GBR IgG, Ctrl IgG or the NMDA receptor–targeting antibody 003-102 IgG (positive control^30^). Following incubation, the medium was replaced with fresh medium containing Alexa Fluor 488–conjugated goat-anti-human secondary antibody (1:200, at saturated conditions), and neurons were incubated for an additional 1 h at 37 °C. Cells were then fixed in 4% PFA for 20 minutes at RT, followed by blocking and permeabilization for 1 h at RT in PBS containing 10% normal goat serum, 10% bovine serum albumin, and 0.1% Triton X-100. Neurons were subsequently incubated with a CF568-conjugated goat-anti-human secondary antibody (Biotium; 1:200) for 2 h at RT. Coverslips were then mounted onto glass slides for imaging.

Dendritic images were acquired on a Zeiss LSM 900 Airyscan confocal microscope (Carl Zeiss Microscopy GmbH, Jena) using a 63×/1.4 NA oil-immersion objective. Imaging parameters were kept constant across all acquisitions (pinhole: 5 AU/264 µm; laser power: 488 nm at 2%, 561 nm at 2%; detector gain: 700 V; pixel size: 0.043 µm × 0.043 µm × 0.150 µm).

Images were analyzed using Imaris x64 version 9.1.4 (Bitplane AG, Oxford Instruments, UK). Reconstructions of neuronal surface–expressed receptors (AF488 signal) and internalized receptors (CF568 signal) were generated using the intensity-based surface function. The ratio of the total reconstructed volume per image for each marker was calculated to quantify receptor internalization.

### Immunostainings and structured illumination microscopy (SIM)

To evaluate the synaptic binding properties of the patient derived GBR IgG to pre- or post-synaptic sites, triple stainings of human IgG and pre- and postsynaptic markers were performed in cultured neurons. At 14 DIV, neurons were incubated for 30 minutes at 37 °C with 10 µg/ml solution of patient derived GBR IgG fractions or Ctrl IgG, then fixed with 4% PFA. Cells were blocked with 10% normal goat serum and 10% bovine albumin serum in PBS, then incubated with the secondary antibody goat anti-human Cy3 (Dianova) 1:200. Subsequently, to evaluate the localization of neuronal bound GBR IgG in the synaptic area, samples where further permeabilized with 0.1% Triton and stained for two additional markers. Therefore, the stained cultures were divided in two subgroups: in one we identified glutamatergic pre- and post-synaptic terminals using rabbit-anti-Homer1 AB 1:500 (Synaptic Systems) and guinea pig-anti-VGlut AB 1:500 (Synaptic Systems); while in the second group, GABAergic synapses were marked using mouse-anti-Gephyrin AB 1:1000 (Synaptic Systems) and guinea pig-anti-VGAT 1:500 AB (Synaptic Systems). The secondary AB used were goat-anti-rabbit and goat-anti-mouse Alexa Fluor 647 (Invitrogen) 1:200 for Homer1 and Gephyrin, respectively, and donkey-anti-guinea pig CF488 (Biotium) 1:200 for VGlut and VGAT. Finally, cells were mounted on glass slides.

Samples were imaged with an Elyra 7 lattice SIM microscope system (Carl Zeiss Microscopy GmbH), using a 63x oil-immersion objective with a NA of 1.4 (Plan-Apochromat 63x/1.4 Oil DIC M27, Carls Zeiss Microscopy GmbH). All recording parameters were kept constant during the experiment (laser power 488 nm at 5.0%, 561 nm at 2.5% and 642 nm at 2.0%; pixel size at 0.03 µm x 0.03 µm x 0.11 µm). Acquisition and rendering of the lattice SIM images were performed in ZEN 3.0 (black SR edition) software by Zeiss. For rendering, a baseline cut was performed, followed by selecting the “best fit” option to reconstruct the super-resolution image from the 15 phases taken per image plane.

For the processing of the SIM images, after rendering, the Imaris x64 9.1.4 (Bitplane AG, Oxford Instruments, UK) software was used. The czi image file was loaded into the software, for each channel the volumetric structure was produced using the surface function, thresholding at the absolute intensity and filtering for the Number of Voxels Img = 1, then the Matlab based coloc-coloc function was used to determine the volumetric structure of colocalization between channels. Volume values were then transferred to an Excel file, where they were computed. The colocalization between the receptor and each marker was expressed as a ratio between the colocalization volume value and the total volume of the corresponding marker.

### STED microscopy

Stimulated emission depletion (STED) imaging was performed similar as described previously ^85^ using an Expert Line Abberior microscope system (Abberior instruments, Göttingen, Germany) with an Olympus (UPlanXApo) 60x oil objective with a numerical aperture of 1.42. The system contained three pulsed fluorescence excitation laser modules for wavelengths of 485, 561, and 640 nm, spectral detection module, and avalanche photodiode (APD) detectors. For STED imaging the system uses a pulsed high-power STED laser module with a wavelength of 775 nm (3 W laser power, repetition rate of 40 MHz). Each image was obtained with sequential line scanning by switching between the needed excitation lasers line-wise during each recording. Each image contained a three channel-recording for the human GBR IgG, the presynaptic active zone marker Bassoon (guinea pig; Synaptic Systems), and the postsynaptic density marker Homer1 (rabbit) for excitatory or Gephyrin (mouse) for inhibitory synapses. All channels were imaged with a dwell time of 5 µs/pixel and a threefold line accumulation. Based on the secondary antibody signals, the excitation laser power for each channel was adjusted to prevent over-or underexposure effects. For GBR IgG, the secondary antibody donkey-anti-human ABBERIOR STAR RED (Abberior) was excited with 640 nm (3.5% laser power) and with a STED laser power of 15% and the detection window was 650 – 755 nm. For Bassoon, the secondary antibody goat-anti-guniea pig ABBERIOR STAR 460L, a long stokes-shift dye, was excited with 440 nm (70.7% laser power) and with a STED laser power of 15% and the detection window was 571 – 630 nm. For Homer1 and Gephyrin, the secondary antibody goat-anti rabbit and goat-anti-mouse ABBERIOR STAR ORANGE (Abberior) were excited with 561 nm (23.6 and 47.2% laser power, respectively) and with a STED laser power of 15% and the detection window was 571 – 630 nm.

### Surgery procedures on mice

#### Osmotic pump and intraventricular catheter implantation

Ten to fourteen weeks old mice, housed one per cage, were implanted with two micro-osmotic pumps (model 1002, Alzet, Durect Corporation, Cupertino, CA) for intraventricular perfusion. As previously reported^86^, osmotic pumps have a volume of 100 µl each, a flow rate of 0.25 µl/h and a duration of perfusion of 14 days. On the day of surgery each pump was loaded with 100 µl of 1 mg/ml solution of patient-derived purified IgG (Ctrl IgG, GBR IgG or GBR-ΔSD IgG) and connected to a bilateral cannula. Meloxicam was orally administered to animals one hour before the surgical procedure, then mice were deeply anesthetized with 1.5 to 2.0% isoflurane/oxygen and placed in a stereotaxic frame (Lab StandardTM, Stoelting, Wood Dale, IL, USA). Cannulas were implanted in the lateral ventricles (coordinates from Bregma: −0.2 mm AP, ±1.0 mm ML; −2.2 mm DV) and the osmotic pumps placed subcutaneously in the back of the mice. Animals were sutured, returned to home cages, and checked daily for the following seven days. During the 14 days of IgG infusion a subset of mice underwent habituation and behavioral tests; after the 2-weeks treatment the mice were sacrificed to perform electrophysiological studies in acute hippocampal brain slices.

#### Implantation of CA3 and CA1 recording electrodes for in vivo LFP recordings and intrahippocampal injections

Seven to three days before surgery animals were handled for at least 10 minutes each, to get used to the experimenter. On the day of surgery, Meloxicam was orally administered one hour before the surgical procedure, then animals were anesthetized with 1.5 to 2.0% isoflurane/oxygen and placed in a stereotaxic frame. The head skin was incised and clamped open to expose the skull. With a dentist driller K.1070 (Foredom, Bethel, USA) holes for ground and fixation screws, electrodes and injections were drilled (coordinates were measured from Bregma, electrodes: CA3 −1.5 mm AP, −1.8 mm ML, −2.0 mm DV; CA1 −2.3 mm AP, −1.6 mm ML, −1.45 mm DV. Injections: CA3 −1.7 mm AP, −3.0 mm ML, −1.7 mm DV; CA1 −2.3 mm AP, −2.9 mm ML, −1.65 mm DV. Fixation screw −3.0 mm AP, +3.0 mm ML. Ground screw −0.5 mm AP, +1.0 mm ML).

First, fixation and ground screws, made with stainless steel uncoated wire (Science Products GmbH, DE), were inserted and glued to the skull with cyanoacrylate glue. Electrodes prepared from platinum/iridium perfluoroalkoxy polymers coated wire (101R-3T, Science Products GmbH, DE) were then lowered in their respective holes and fixed in the same way. After that, dental cement Paladur (Kulzer GmbH, DE) was used to hold the electrodes and prepare a base to hold a 4-pin holder, with extra care to keep the injection holes open. The pin-holder was lowered, the electrodes and ground inserted in the corresponding holes, and everything was fixed with dental cement to create a “dummy head”. After the dental cement dried, the skin was sutured. At the end of surgery, the animal was returned to the home cage, kept under observation, and treated with Meloxicam for the following three days after surgery. On day six after surgery, animals were brought again to the surgical frame to perform intrahippocampal injections. The skin was reopened to expose the open holes for injection. With a nano injector Micro 2T (WPI, Saratosa, USA), a stereotaxic arm inclined at 45° and a very thin injection pipette (tip opening around 10μm) pulled from borosilicate glass, 1 µl per injection hole of 1 mg/ml patient-derived purified IgG (GBR IgG or Ctrl IgG) was injected at a 4 nl/s rate. Animals were re-sutured, and connected to the tethered system in the recording cage. Video acquisition and the local field potential (LFP) recordings were conducted for the following 24 h.

### Behavioral Tests

#### Novel Object Recognition test

After recovery from osmotic pump implantation surgery, animals were habituated for five days before the test for 5−10 minutes each day in a 41x41x32 cm open field arena.

The NOR was performed to assess memory, based on the innate preference of rodents to explore novel items. All tests were carried as previously described^87^ by an investigator blinded to the animals’ treatment group.

Animals were divided in two independent cohorts. The first cohort was used to assess memory formation using the NOR paradigm with a 3 h retention interval. The second cohort was used to evaluate the effect of patient-derived IgG preparations on memory retrieval using the NOR paradigm with a 24 h retention interval. To test the initial memory formation in this cohort using a different test paradigm, animals were additionally tested using the NOL paradigm within a 3 h retention interval.

On the test day, mice were placed in the arena with two identical objects for the familiarization phase. A video recording of the animals exploring the objects was acquired for a total time of 10 minutes. After a retention time of either 3 h or 24 h in the home cages, mice were placed once again in the open field arena, where one of the familiar objects was replaced by a novel object and positioned in a novel location within the box. Which of the two familiar object was replaced with the novel one, and in which position the novel one was placed, was randomized among trials. The explorative behavior towards the familiar and novel objects was evaluated for 10 minutes video recordings and then manually analyzed.

Animals were tested on day 7 post-surgery (3 h retention time) or on day 6/7 post-surgery (24 h retention time).

#### Novel Object Location test

The NOL was carried as the previously described NOR, with the only difference that one of the familiarized objects was moved to a different location within the open field arena during the test phase. Animals were tested only with 3 h retention time and their explorative behavior was evaluated for 10 minutes on day 7 post surgery.

#### Analysis of behavioral tests

For the analysis of behavioral tests (NOR and NOL), an in house-coded software was used to evaluate the time spent by each animal exploring the objects. All analyses were performed by a blinded investigator unaware of treatment groups. Nose poke or orientation of the nose to the object in a range of 2 cm from the object itself was considered an explorative behavior and timed. The explorative times (in [s]) were indexed by a preference score (PS), based on the time spent exploring the novel object or location (T_NO_) and the familiar object (T_FO_): *_PS_* = 100 ∗ *_TNNN_*/ (*_TNNN_* + *_TFNN_*) . Animals that did not reach a total cumulative exploration time of at least 20 seconds per trial, or animals that showed preference for one of the two identical objects during the familiarization phase, were excluded from the analysis.

### Electrophysiological recordings

#### Recordings of primary murine cell cultures

Primary cell cultures hippocampal neurons were used for whole-cell patch-clamp recordings after 11 to 18 DIV. 24 hours before the recording the cell cultures were incubated with either GBR IgG or Ctrl IgG (20 µg/ml), diluted in neurobasal medium. Recording electrodes were pulled from thick-walled borosilicate glass (Science products, Hofheim, Germany) and filled with intracellular recording solution containing (in mM) 135 CsMeSO_4_, 3 CsCl, 10 HEPES, 0.1 spermine, 2 Mg-ATP, 0.3 Na-GTP, 10 phosphocreatine, 0.2 EGTA, 2 QX-314 bromide (Tocris, Bristol, UK), adjusted to pH 7.3 with 1N CsOH, osmolality 280 mOsmol/kg, and had a final pipette resistance of 3 – 5 MΩ. For evaluation of AMPAR-mediated sEPSC voltage-clamp recordings were performed for 100 s at a holding potential of −70 mV in aCSF-1 containing (in mM) 125 NaCl, 25 NaHCO_3_, 25 glucose, 2.5 KCl, 1.25 NaH_2_PO_4_, 1 MgCl_2_, 2 CaCl_2_; saturated with 95% O_2_ and 5% CO_2_, pH 7.3, osmolality between 300 to 310 mOsmol/kg. In about half of the experiments 5 µM baclofen was added to the aCSF-1 and perfused for at least 5 minutes to systemically activate GABA_B_Rs before repeating the sEPSC recordings. Traces of sEPSC recordings were low-pass filtered at 1 kHz (Butterworth Low-Pass) and analyzed with Mini Analysis software (Synaptosoft, Decatur, GA, USA) using AMPAR-mediated sEPSC compatible templates. The automatically detected events were then visually examined and erroneous events were removed from further analysis.

#### Acute slice preparation and recording solutions

15 to 20 days after pump implantation, acute hippocampal brain slices were prepared as described previously^88^. In short, after decapitation of mice under deep Isoflurane anesthesia, the brain was quickly removed from the skull and placed in ice-cold (4 °C) aCSF. For whole-cell patch-clamp and fEPSP recordings in CA1, the preparation and slicing were performed in ice-cold (4 °C) protective aCSF (paCSF) composed of (in mM): 95 N-methyl-D-glutamine, 30 NaHCO_3_, 20 HEPES, 25 glucose, 2.5 KCl, 1.25 NaH_2_PO_4_, 2 thiourea, 5 sodium ascorbate, 3 sodium pyruvate, 10 MgSO_4_, 0.5 CaCl_2_, 12 N-acetylcysteine, adjusted to pH 7.3 and an osmolality 300 – 310 mOsmol/kg, saturated with 95% O_2_ / 5% CO_2_. Tissue blocks containing the hippocampal formation were mounted on a vibratome (Leica, Wetzlar, Germany; Leica VT1200 S) and coronal slices were cut at 350 µm thickness. Slices were then incubated in the same paCSF at 34 °C for 12 minutes.

For whole-cell patch-clamp recordings, slices were transferred to an incubation beaker with protective aCSF-2 (in mM: 125 NaCl, 25 NaHCO_3_, 25 glucose, 2.5 KCl, 1.25 NaH_2_PO_4_, 1 MgCl_2_, 2 CaCl_2_, 2 thiourea, 5 sodium ascorbate, 3 sodium pyruvate, 12 N-acetylcysteine, adjusted to pH 7.3 osmolality of ∼300 mOsmol/kg, equilibrated with 95% O_2_ / 5% CO_2_) and stored at room temperature for at least one hour or until recording. For subsequent recordings of fEPSP and LTP in the CA1, the slices were transferred to a submerged incubation beaker filled with aCSF-3 (in mM: 124 NaCl, 26 NaHCO_3_, 10 glucose, 3.4 KCl, 1.2 NaH_2_PO_4_, 2 MgSO_4_, 2 CaCl_2_; equilibrated with 95% O_2_/5% CO_2_, pH 7.3, osmolality 300 – 310 mOsmol/kg) and stored at room temperature for at least one hour prior recording.

Slices for the fEPSP recording in the CA3 upon MF stimulation were prepared in ice-cold (4 °C) sucrose-based saCSF containing (in mM): 50 NaCl, 25 NaHCO_3_, 10 glucose, 2.5 KCl, 1 NaH_2_PO_4_, 7 MgCl_2_, 0.5 CaCl_2_, and 150 sucrose, saturated with 95% O_2_ and 5% CO_2_. Sagittal slices were cut at 300 μm thickness and incubated in saCSF at 35°C for 30 minutes. Before the recording, slices were transferred into a submerged incubation beaker filled with aCSF-4 containing (in mM): 119 NaCl, 26 NaHCO_3_, 10 glucose, 2.5 KCl, 1 NaH_2_PO_4_, 1.3 MgCl_2_, and 2.5 CaCl_2_, saturated with 95% O_2_ and 5% CO_2_ and stored at least 1 h at RT prior recording.

#### Whole-cell patch-clamp recordings of CA1 pyramidal neurons

For whole-cell patch-clamp recordings slices were placed in a submerged recording chamber on an upright microscope (Carl Zeiss Microscopy GmbH), perfused with aCSF-1 (see above) and cells were visualized using difference-interference (DIC) optics with an 63x water-immersion objective (Carl Zeiss Microscopy GmbH). Electrophysiological recordings were conducted with a HEKA EPC-10/2 amplifier (HEKA Elektronik, Lambrecht/Pfalz, Germany) with a sampling frequency of 20 kHz (postsynaptic currents) or 100 kHz (action potentials) and were low-pass filtered at 2.9 or 10 kHz with an implemented Bessel filter of the amplifier or with a MultiClamp 700B amplifier (Molecular Devices, Sunnyvale, CA, USA) and digitized using an Axon Digidata 1550B digitizer and Clampex 11.2 software (Molecular Devices, Sunnyvale, CA, USA). Somatic recordings were performed in pyramidal neurons in the CA1 and CA3 area of the hippocampus using pipettes pulled from thick-walled borosilicate glass (Science Products, Hofheim, Germany; Flaming/Brown Micropipette Puller Modell P-1000, Sutter Instrument, CA, USA) to resistances between 3−8 MΩ when filled with an internal solution according to the respective protocol.

To determine subthreshold membrane properties (input resistance, membrane time constant, subthreshold resonance; **Fig. 4a-b** and **Extended Data Fig. 4**), recordings were performed in the continuous presence of baclofen (1 µM). Pipettes were filled with an intracellular solution containing (in mM): 120 K-gluconate, 20 KCl, 10 Hepes, 0.1 EGTA, 4 Mg−ATP, 0.2 Na_2_−GTP, 10 Na_2_−Phosphocreatine; pH 7.3 (adjusted with 1N KOH), osmolality 290 mOsmol/kg. A series of current steps ranging from −100 pA to 80 pA in 20 pA increments was applied to the soma and the voltage responses recorded in current clamp mode. Input resistance (R_in_) and membrane time constant (τ_m_) were derived from subthreshold current steps that produced voltage deflections within ±10 mV. R_in_ was calculated as the slope of a linear fit to the voltage-current relationship. τ_m_ was computed by fitting a single exponential to the averaged post-pulse voltage relaxation starting 2 ms after pulse termination.

To determine membrane resonance properties of CA1 and CA3 pyramidal cells, an impedance amplitude profile (ZAP) protocol was applied. A chirp current of constant peak-to-peak amplitude (40 pA) and linearly increasing frequency from 0 to 60 Hz over 90 s was injected into the soma. Measurements were obtained at two membrane potentials close to −60 and −75 mV. The protocol consisted of three consecutive sweeps and was repeated at both holding membrane potentials for as long as stable whole-cell recordings could be maintained. Voltage and current traces were preprocessed to remove spikes and baseline drift. Spikes were identified as rapid depolarizations exceeding −40 mV. Traces were high-pass filtered at 0.01 Hz after mirror-padding before the start of the ZAP stimulus to remove slow voltage drifts.

Current and voltage signals were subjected to Fast Fourier Transform (FFT). Impedance was defined as Z(f) = FFT(V)/FFT(I). Impedance amplitude at frequency f was defined as the absolute value of Z(f) and denoted as |Z(f)|. Peak resonance frequency was defined as the input frequency corresponding to the maximum impedance. Mean |Z(f)| was computed within defined frequency bands (theta: 4−10 Hz; gamma: 30−60 Hz) for each pyramidal cell.

For measurements of GABA_A_-receptor eIPSCs, an internal solution containing (in mM) 125 Cs-methanosulfonate, 7 NaCl, 1 HEPES, 0.1 EGTA, 2 MgCl2, 0.1 CaCl, 4 Mg−ATP, 0.3 Na_3_−GTP, 5 Na_2_−Phosphocreatine; pH 7.3 (adjusted with CsOH), osmolality 290 mOsmol/kg, was used. For recordings of AMPA-receptor eEPSCs as well as GIRK-dependent slow hyperpolarizations (sIPSPs) and APs, Cs-methanosulfonate was replaced by K-methanosulfonate, which enabled the measurement of changed membrane potentials in current-clamp. For voltage-clamp recordings, cell membranes were clamped at a −70 mV holding potential, current-clamp recordings were performed at the resting membrane potential of the neuron. The series resistance was compensated (60 – 70%) and checked regularly during the recording.

To investigate the influence of GBR IgG onto the glutamatergic synaptic transmission, EPSCs were evoked by electrical stimulation (Isoflex, A.M.P.I., Jerusalem, Israel) of the SC in SR with an aCSF-1 filled bipolar theta-glass electrode placed in a minimal distance of 200 µm to the recorded neuron to avoid direct activation. With 200 µs impulses and a step-wise increase of the electrical stimulation between 50 and 400 µA, an input-output-curve of eEPSCs was recorded to determine the half-maximal stimulation strength (Max/2) for each individual cell. This Max/2 stimulation was utilized for a paired-pulse protocol with an ISI of 50 ms to measure short-term plasticity. To isolate eEPSCs pharmacologically, 50 µM (2R)-amino-5-phosphonovaleric acid (AP-5, #0106; Tocris Bioscience, Ellisville, MO, USA) and 20 µM bicuculline (#0109; Tocris, Bioscience, Ellisville, MO, USA) were added to the perfused aCSF-1 that inhibit NMDA- and GABA_A_-receptors, respectively. In order to evaluate the pathological effect of GBR IgG on the activability of GABA_B_ heteroreceptors, the eEPSC measurements were repeated under perfusion of the agonist (±)-baclofen (5 µM; #B5399, Sigma) and subsequently with perfusion of the potent GABA_B_R antagonist CGP55845 hydrochloride (10 µM; #1248, Tocris).

The activation of GABAergic interneurons and the resulting activity of GABA_B_-autoreceptors in presynaptic terminals was investigated by electrical stimulation in the SLM. This layer contains cholecystokinin (CCK) positive interneurons and neurogliaform cells (NGF), both of which express GABA_B_Rs in high density at their axon terminals^89,90^. These were excited by short trains of 10 impulses in a frequency of 5 Hz (interstimulus interval 200 ms). The resulting eIPSCs were isolated by blocking ionotropic glutamate receptors (50 µM AP-5 for NMDA, 10 µM 6-cyano-7-nitroquinoxaline-2,3-dione [CNQX] for AMPA) and by clamping the cells at a holding potential of −40 mV increasing the driving force of Cl^−^ ions. For each cell, the eIPSC trains were recorded in 10-fold repetition before and after bath perfusion of 10 µM CGP55845 in aCSF-1.

To characterize postsynaptic GABA_B_Rs, the SLM interneurons were stimulated with short 100 Hz trains of 5 impulses, which induces GABA release sufficient to diffuse to the postsynaptic membrane, where it binds to and activates GABA_B_Rs. Given the high frequency stimulation, modulation of the slow presynaptic autoreceptors via feedback inhibition is not expected, thereby allowing for a decoupled examination of the postsynaptic GABA_B_R. With this protocol the GIRK channel mediated sIPSPs were recorded in current-clamp at resting potential and under pharmacological blockade of AMPA-, NMDA-, and GABA_A_-receptors (see above). The effects on intrinsic excitability were investigated with a continuously increasing current injection of 0 – 300 pA within 200 ms leading to a ramp depolarization (slope 1.5 mV/s) that initiates APs. This was done without and with a preceding 100 Hz burst stimulation for the postsynaptic GABA_B_R activation and the effect on the latency to the first AP and the number of APs were analyzed. To further investigate the ability of GABA_B_R activity to negatively modulate the AP generation, we used a protocol previously described as spike blocking efficacy^35^. For that, one stimulation electrode was placed in the SC to induce a single AP by applying an extracellular stimulation in an intensity strong enough for a spiking rate of ∼90−95% (20 sweeps). With a second stimulation electrode placed in the SLM, high-frequency stimulations (HFS) of 10 impulses at 100 Hz are applied 250 ms prior to the SC stimulation to activate the GABA_B_R-mediated inhibition. CA1 pyramidal cells were recorded 1) in aCSF-1 and SC stimulation alone (−HFS), 2) with preceding 100 Hz train (+HFS) for 20 sweeps each. Most of the experiments were repeated after bath perfusion of 10 µM CGP55845 with 3) −HFS and 4) +HFS. The spiking probability was determined by dividing the number of successfully initiated APs by the total number of sweeps.

#### fEPSP recording and LTP induction

Recordings of fEPSP in the hippocampal CA3 upon stimulation of the mossy fiber (MF) pathway, were done as previously described^91^. In brief, the recordings were performed at 21 – 24 °C in a submerged recording chamber perfused with aCSF-4. MFs were stimulated at 0.05 Hz using a low-resistance glass electrode filled with aCSF-4, which was placed close to the internal side of granule cell layer of the dentate gyrus. The recording electrode filled with aCSF-4 was placed in *stratum lucidum* of area CA3. The MF origin of fEPSPs was verified by the pronounced facilitation of synaptic response amplitudes upon 1 Hz stimulation (not shown) or paired pulse stimulation at 25 Hz. PPR was calculated by dividing the amplitude of the second pulse with the amplitude of the first pulse. Means of both groups were used for statistical comparisons. Example traces represent average of triplet recording. In addition, we confirmed that fEPSPs were generated specifically by MFs by utilization of the type II metabotropic glutamate receptor agonists LCCG1 ((2S,1′S,2′S)-2-(carboxycyclopropyl) glycine), 10 µM) during the recording or DCG IV ((2S,2’R,3’R)-2-(2’,3’-dicarboxy-cyclopropyl) glycine; 1 µM) at the end of each recording (not shown). These agonists specifically suppress transmitter release from MF terminals in CA3, but not from neighboring associational-commissural fiber synapses^92^. Only responses that were inhibited by >75% were accepted as MF signals (28 slices excluded with DCG IV and 20 slices with LCCG1). Input-output (I/O) relationship was determined by applying at least three different stimulation intensities per slice and measuring the amplitudes of presynaptic fiber volley (FV) and postsynaptic fEPSP amplitude. Each intensity was recorded in triplets and the average was calculated. If it was not possible to stimulate with three different intensities, the slice was not included in the analysis (Ctrl IgG: n=23 slices, GBR IgG: n=21 slices included).

For fEPSP recordings in the hippocampal CA1, slices were placed in an interface chamber (BSC2, Scientific System Design Inc), which allowed for maintaining the temperature at 35 °C, and continuously perfused with aCSF-3 at 1.5 to 2.5 ml/min flow rate through a peristaltic pump (ISM831C, ISMATEC). fEPSP were recorded from the SR of the CA1 region using a glass microelectrode pipette filled with aCSF-3 (tip resistance 3 to 7 MΩ). The potentials were elicited every 30 s with a custom-made platinum-iridium bipolar electrode placed along the SC fibers pathway and connected to an Iso-Flex stimulator (Iso-Flex, Science Products GmbH). Data acquisition and stimulation protocols were performed with a HEKA EPC-10 amplifier (HEKA EPC-10, HEKA Elektronik Dr. Schulze GmbH). Data were acquired at a 20 kHz sampling rate, filtered at 2.9 kHz with a Bessel filter, and digitized.

Before the LTP induction protocol, input-output (I/O) curves with increasing stimulation intensities ranging from 0 mA to 1 mA in steps of 0.1 mA were obtained. This allowed to establish the half-maximal response stimulation intensity then used for the baseline test pulse and the LTP induction protocol. After recording 20 minutes of stable test pulses baseline, LTP was induced with a TBS, comprising 20 bursts, each consisting of 4 pulses, at an inter-burst frequency of 5 Hz and an intra-burst frequency of 100 Hz^93^. The protocol was repeated in seven sweeps with an interval of 30 s. Responses to test pulse were recorded thereafter every 30 s for 60 minutes to measure LTP. In a subset experiment, after the initial I/O curve and the first 20 minutes test pulse baseline, an aCSF-3 containing 10 µM CGP55845 (Tocris Bioscience, UK) was perfused to block GABA_B_Rs. During the CGP55845 perfusion, an additional test pulse baseline was recorded, thereafter the LTP was induced and the post induction fEPSP recorded for 60 minutes.

For the analysis of the fEPSPs, the slope measure of each potential (calculated as mV over ms) was evaluated from 20% to 80% of the rise time with the NeuroMatic plugin of IGOR Pro 6.37 software (WaveMetrics, Lake Oswego, OR). Erroneous traces (high noise, spontaneous inputs interfering with the evoked signal) were discarded manually before analysis. The values of slopes obtained were then computed from raw data for the I/O curve or from normalized slopes to the median of the last 10 minutes of baseline for the LTP experiments.

### *In vivo* LFP recordings and analysis

#### Recording configuration

Animals after surgeries were equipped with a head-fixed 4-pin holder, providing the possibility to connect them to homemade cables and record LFP signals from the CA3 and CA1 areas of the hippocampus. To record the LFP, signals from intracerebral electrodes were transmitted to the pre-amplifier and then to amplifiers EPMS-07 (NPI electronics GmbH, DE), amplified at gain 10, filtered with a highpass filter 0.3 Hz and a lowpass filter 1.7 kHz, and digitized by a PowerLab 16/35 digitizer (ADInstruments, Oxford, UK). Sample frequency of 4 kHz was set in the acquisition software, LabChart Pro 8.1.28 (ADInstruments, Oxford, UK).

The LabChart software allowed the possibility to record up to 16 channels at the same time, and to synchronize LFP recordings with a video recording of the animals’ behavior, coming from a home surveillance style system composed of infra-red cameras THC-T140 connected to a digital video recorder DVR-204Q-K1 (HiLook, CHN) for day and night recordings. The whole system was set to record at the same time from four plexiglass 21x21x28 cm homemade cages to which animals were habituated for five days after surgery to familiarize themselves with the new environment and get used to moving around with the weight of the connected tethering cable. On day six after surgery, after the intrahippocampal injection of purified patient IgG fractions, animals were placed in the system and behavioral videos paired with LFP traces were recorded for 24 h.

#### Preprocessing of signals

The simultaneous recordings of the extracellular signals from CA1 and CA3 areas in freely moving mice were described above. The LFP signals were processed using custom-written code in MATLAB 2023a (MathWorks). Movement artifacts were detected in a two-step, iterative process. First, a threshold of 10 times the standard deviation (SD) of each signal was used to detect large-amplitude artifacts. After setting these periods to NaN (missing value), the SD was recomputed, and a final threshold of 7xSD was applied. The detected artifacts were then visually inspected and confirmed against simultaneously recorded cage movies. To ensure robust exclusion of artifactual data, the verified artifact periods from both CA1 and CA3 signals were concatenated and then extended bidirectionally by 2 seconds. This concatenation was a required step to maintain a consistent time base for the interareal coupling analysis. The resulting signals had a duration of 157.05 ± 27.73 minutes in the Ctrl IgG group (n=10 mice) and 151.83 ± 26.49 minutes in the GBR IgG group (n=9 mice). The signals were downsampled to 250 Hz, preceded by a low-pass anti-aliasing FIR filter with a cutoff of 100 Hz and a narrow transition (*lowpass* function, steepness = 0.99).

#### Phase-amplitude coupling (PAC) analysis

For each mouse, we quantified the theta-gamma phase-amplitude coupling between CA3 and CA1 using a well-established modulation index (MI) based on the Kullback-Leibler distance (KL-distance)^45,46^; the main Matlab code is available in https://github.com/tortlab/phase-amplitude-coupling. The continuous LFP signal was first segmented into 1-minute overlapping time bins (30-second step; e.g. see^94,95^). This bin length was chosen to provide a stable estimate of the coupling^45^. For each time bin, the instantaneous phase and amplitude time series were extracted (*Hilbert* function). This process was applied to multiple, narrowly filtered frequency-band pairs, with the phase-providing signal filtered from 5 to 12 Hz in steps of 1 Hz (8 theta bands, CA3) and the amplitude-providing signal filtered from 30 to 90 Hz in steps of 5 Hz (13 gamma bands, CA1) (*eegfilt.m*, EEGLAB toolbox^96^). For each of these 104 frequency-band pairs (in a 13x8 matrix), a phase-amplitude distribution was constructed by binning the theta phases into eighteen 20° intervals and calculating the mean gamma amplitude in each phase bin. This distribution was normalized in each phase bin by the sum of amplitude values across bins, in order to create a probability density-like function. The MI was then calculated as proportional to the KL-distance between this observed distribution and a uniform distribution (the no-coupling case). The higher the MI (as inferred by KL-distance), the further away the phase-amplitude distribution is from the uniform distribution.

To assess statistical significance, the empirical MI of each bin was compared to a distribution of 200 surrogate MI values. Each surrogate was generated by applying a random circular shift to the gamma amplitude envelope while leaving the theta phase time series intact, followed by re-computing the coupling^45^. This shuffling procedure disrupts the specific temporal relationship between the two signals while preserving the autocorrelation and spectral properties of the original time series. If an empirical MI was larger than the 99^th^ percentile of its bin-specific surrogate distribution, the time bin was considered statistically significant. The mean and SD of the surrogate distribution were further used to z-score the empirical MI to achieve a normalized measure of coupling strength. Any frequency pair in a given bin with a non-significant MI was set to NaN.

#### Comodulogram and coupling parameter extraction

For each mouse, a single, summary comodulogram^45,46^ was computed by taking the mean of the z-scored MI matrices from all bins containing at least one significant frequency pair. In this averaging process, any non-significant frequency pair (previously set to NaN) was discarded, ensuring the final average reflects the strength of only the significant coupling events.

To extract quantitative parameters, this summary comodulogram was first lightly smoothed with a 2D Gaussian filter (sigma = 0.5). From this smoothed comodulogram, we extracted the peak MI and its corresponding peak theta and gamma frequencies on a per-mouse basis. To quantify spectral specificity, each mouse’s hotspot in this map was defined as the contiguous region where the MI exceeded 75% of the peak value. The coupling area was then calculated for each mouse as the number of frequency pairs within this hotspot region, and the coupling theta and gamma bandwidths were defined as the frequency ranges of this region in x- and y-axes, respectively. Separately, to define a standardized and unbiased set of frequency bands for the subsequent spectral analysis, a region of interest (ROI) was created from the grand average comodulogram of the Ctrl IgG group. This data-driven ROI defined the canonical frequency bands for strong coupling as 6–10 Hz for theta and 50–85 Hz for high-gamma.

To specifically test if GBR IgG differentially affected low-versus high-gamma coupling with theta, we first created a gamma-coupling profile for each mouse. This was achieved by averaging the z-scored MI values across the theta band (6−10 Hz), resulting in a single vector representing MI strength as a function of gamma frequency. From this profile, we then calculated the mean MI within the low-gamma band (30−50 Hz) and the high-gamma band (50−85 Hz). These two values for each mouse were then used for statistical comparison between the groups.

#### Power Spectral Analysis

Power spectral density (PSD) was calculated for each individual 1-minute time bin using Welch’s method (*pwelch* function, 2-second Hamming windows with 50% overlap). To control for potential variability in recording impedance and to provide a measure of the spectral profile, all analyses were performed on relative power.

To directly test the power dynamics that potentially support hyper-coupling, we performed a coupling-conditional analysis. First, the single frequency pair corresponding to the peak of the grand average comodulogram in Ctrl IgG (8 Hz theta, 70 Hz gamma) was considered as the canonical communication channel, in the present data. Then, for each mouse, we selected only the PSDs from the time bins where the MI at this specific frequency pair was statistically significant. A single, grand average conditional PSD for each mouse was then calculated by first averaging these selected PSDs. This final mean PSD was then normalized by its total power, integrated over a frequency range of 1 to 125 Hz. This final relative spectrum was used to calculate the average power for our key frequency bands, namely in the data-driven theta (6–10 Hz) and high-gamma (50–85 Hz) bands defined above as well as a low-gamma band (30–50 Hz), and the resulting values were log10-transformed for statistical testing.

To compute the overall relationship between CA1 and CA3 power spectra, we calculated the ratio of the CA1 over CA3 PSD at each frequency. For each mouse, this ratio spectrum was summarized by its median value, which was then converted to dB units (*10*log10).

### Computational neural network model

#### Model description

To mechanistically link the altered synaptic kinetics to the abnormal CA1 network oscillations under GBR IgG, we employed a well-established, biophysical neuronal network model of CA1^54–57^. This representative model largely captures the core excitatory-inhibitory (E-I) interactions thought to underlie the hippocampal theta-nested gamma oscillations^39,40,46,56^. Specifically, it incorporates three key Hodgkin-Huxley-type neuronal populations: pyramidal cells (‘E-cells’) and fast-spiking PV^+^ cells (‘I-cells’), which form the gamma-generating subnetwork, and oriens lacunosum-moleculare interneurons (O-LM or ‘O-cells’), which preferentially fire in the theta band. To ensure computational efficiency, the model utilizes a reduced network architecture consisting of 10 cells per population. To maintain the characteristic oscillatory dynamics of larger-scale biological networks, synaptic weights are scaled to represent the collective drive of larger ensembles (see below), as previously implemented in this model architecture^55–58^. Based on this canonical architecture, the model has been used extensively to explain and predict the emergence of complex LFP dynamics across various behavioral or pharmacological conditions^53,56–58^. As such, it provides a robust, validated framework for investigating how cellular and synaptic dysfunctions scale to pathological population-level rhythms, regardless of the specific absolute frequency bands involved.

In this model, in addition to within-CA1 synaptic inputs, the E- and I-cells receive an external constant excitatory drive (with additive noise) representing SC input; denoted as *ε_E_* (E-drive) and *ε_I_* (I-drive). Parameter values were primarily based on previous studies^56,57^, with the baseline configuration representing the control condition. To simulate GABA_B_R-AB effects, we re-parametrized the Control model by incorporating our experimental observations of: (1) GABA_B_ autoreceptors dysfunction, manifested as an increase in the steady-state IPSC amplitudes onto pyramidal cells (**Fig. 3b**), and (2) GABA_B_ heteroreceptors dysfunction, manifested as a steeper slope of fEPSPs in CA1/SR (**Fig. 5f**).

Because the model primarily simulates persistent network oscillations, we assumed that GABA_B_Rs operate at a steady state during these ongoing rhythms. We therefore mapped the observed synaptic alterations into the model by applying a relative increase in the inhibitory synaptic weights of I- and O-cells onto E-cells (*G_IE_* and *G_OE_*) and a relative increase in the E-drive (*ε_E_*). This approach effectively emulates the functional consequences of GABA_B_R-AB on presynaptic kinetics without requiring the explicit implementation of GABA_B_R molecular signaling. Consequently, while the model is not intended to account for the transient initiation phases of network oscillations, it provides powerful mechanistic insights into the core effects of GABA_B_R-AB on the network hypersynchrony observed in our data. In the following sections, we detail the neuronal and synaptic models, LFP simulations, and the numerical procedures used for setting the numeric and random aspects of the model.

#### Neuron models

E cells: For the pyramidal cells, we use Olufsen et al.^97^ model, as a variation of Ermentrout et al.^98^ model:

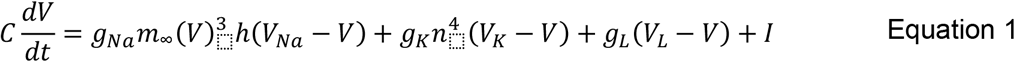

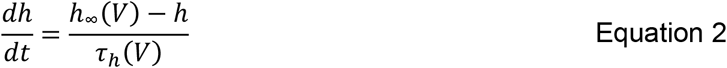

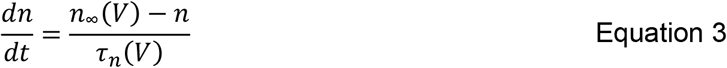

With

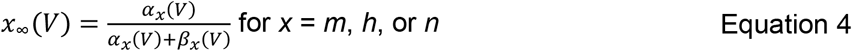

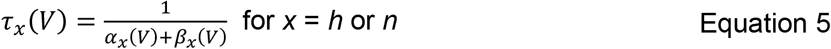

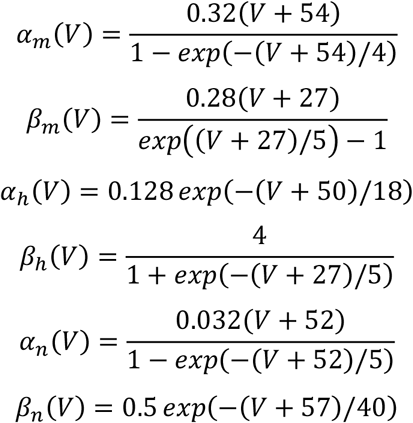

In Equations 1-3, the letters *C*, *V*, *t* and τ, *g*, and *I* denote capacitance density, voltage, time, conductance density, and current density, respectively. The units that we use for these quantities are µF/cm^2^, mV, ms, mS/cm^2^, and µA/cm^2^. For brevity, units will usually be omitted hereafter. The parameter values of the model are *C* = 1, *g_Na_* = 100, *g_K_* = 80, *g_L_* = 0.1, *V_Na_* = 50, *V_K_* = –100, and *V_L_* = –67.

I cells: For fast-spiking PV+ interneurons, we use the model described by Wang and Buzsáki et al. ^99^. Equations 1-4 are the same as in the E cell model, but the Equation 5 is replaced by:

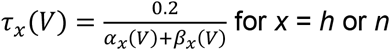

with the rate functions *α_x_* and *β_x_* (*x* = *m*, *h*, and *n*) are defined as follows:

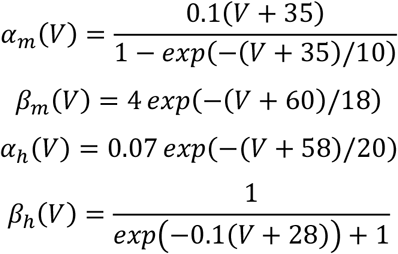

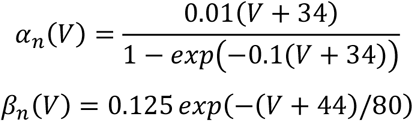

The parameter values, using the same units as for the pyramidal cell, are *C* = 1, *g_Na_* = 35, *g_K_* = 9, *g_L_* = 0.1, *V_Na_* = 55, *V_K_* = −90, and *V_L_* = −65.

O cells: For the oriens lacunosum-moleculare interneurons, we use the model described in by Tort et al.^55^ which is a reduction of the multi-compartmental model described by Saraga et al.^100^ The current-balance equation is given by:

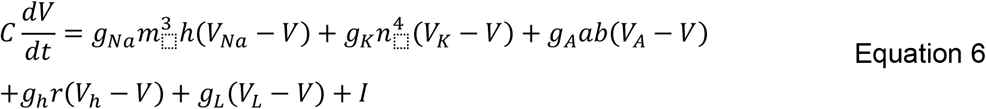

With

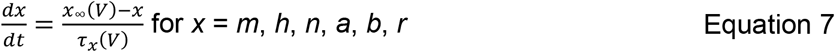

For *x* = *m*, *n*, *h*, the functions *_x∞_*(*_V_*) and *_τx_*(*_V_*) are the same as in Equations 4 and 5, and the rate functions *α_x_* and *β_x_* are defined as follows:

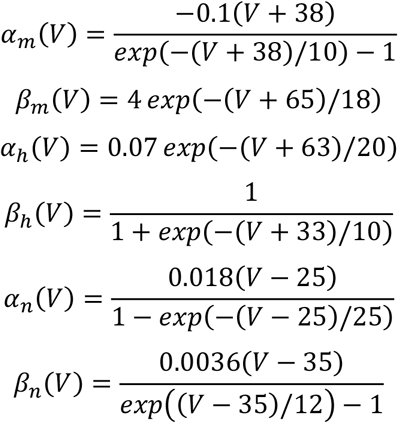

For *x* = *a*, *b*, *r*, the functions *_x∞_*(*_V_*) and *_τx_*(*_V_*) are defined as:

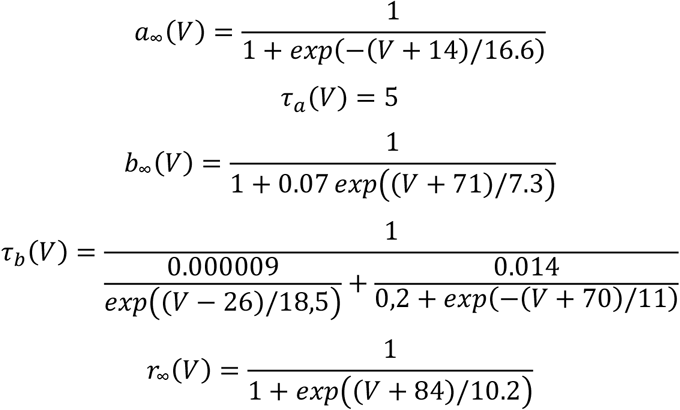

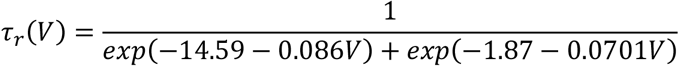

The parameter values, using the same units as for the pyramidal cell, are *C* = 1.3, *g_L_* = 0.05, *g_Na_* = 30, *g_K_* = 23, *g_A_* = 16, *g_h_* = 8, *V_Na_* = 90, *V_K_* = −100, *V_A_* = −90, *V_h_* = −32, *V_L_* = −70 (see also Wulff et al.^57^).

#### Synaptic model

Following previous studies^56,57,98^, we model each synaptic input as a current of the form *I_syn,XY_ = G_XY_/N_X_ s(V − V_syn_)*, where X and Y denote the type of the pre and postsynaptic cell, respectively (i.e. X and Y belong to E, I, or O), *G_XY_* is the maximal synaptic conductance, *N_X_* is the number of *X* cells, *V* is the membrane potential of the postsynaptic cell, and *V_syn_* is the reversal potential. The variable *s* is a normalized double-exponential function characterized by rise (*_τX_*_,*r*_) and decay (*_τX_*_,*d*_) time constants. The synapses were implemented using the *Exp2Syn()* built-in function of NEURON. The IPSPs originating from the O cells synapses were slower than IPSPs originating from the I cells synapses; following these studies, we used *_τE_*_,*r*_ = 0.1 ms and *_τE_*_,*d*_ = 3 ms for E synapses, *_τI_*_,*r*_ = 0.3 ms and *_τI_*_,*d*_ = 9 ms for I synapses, and *_τI_*_,*r*_ = 0.2 ms and *_τI_*_,*d*_ = 20 ms for O synapses. The reversal potential was set to 0 mV for E synapses and to −80 mV for I and O synapses. We used *G_II_* = 0.05, *G_OI_* = 0.2, *G_IO_* = 0.5, *G_IE_* = 0.1, *G_OE_* = 0.15, *G_EI_* = 0.1, *G_EO_* = 0.01 mS/cm^2^. In our model, in addition to the synaptic inputs (*I_syn,XY_*) from X populations within CA1, neurons receive external drive currents representing inputs from upstream regions such as CA3 (via SC) or the entorhinal cortex (via the temporoammonic pathway). We model this drive as a constant (i.e. mean) level of *_εX_* with an additive white noise component. Baseline drives for the Control condition were set to *_εE_* = 0.8, *_εI_* = 0.8, and *_εNN_* = −3.0 µS/cm^2^. Using these biologically grounded values consistent with previous studies^55–57^, the Control model generated simLFPs with theta and gamma peak frequencies at ∼6 Hz and ∼40 Hz, respectively. As noted above, this model is designed to simulate representative CA1 network oscillations. Rather than fine-tuning parameters to precisely match specific experimental frequency bands or absolute peak powers observed in our data, we utilized this established framework to provide mechanistic insights into how synaptic dysfunctions drive network-level hypersynchrony.

#### Model local field potential

The simulated local field potential (simLFP) of the model consisted of the membrane potential of a “passive” E cell programmed inside the network^53,55–57^. This cell receives the same synaptic inputs as the “active” E cells. However, this cell does not send any synaptic output onto other cells and, in addition, it also does not spike given that its external drive current is set to zero. Power spectra of the simLFP were obtained using the Welch’s method (MATLAB *pwelch()* function) with 2-s Hamming windows, 75% overlap.

#### Numeric and random aspects of model

All simulations were carried out using NEURON simulation program version 8.2 (https://www.neuron.yale.edu/neuron/)^101^. The model is publicly available code in ModelDB github page (https://github.com/ModelDBRepository; repository number: 138421), and was simulated with a time step of 25 µs. As initial conditions, the membrane potential of each cell was uniformly distributed between −85 and −60 mV and the channel-gating variables were set to their corresponding steady-state values. Each cell was further randomized using a clamping current (*I_Clamp_*) of random uniform magnitude and random uniform duration between 0 and *t_syn_*/2, where *t_syn_* = 0.5 s is the time when synapses were turned on. In addition to the drive current (see previous section), each cell in the network also received additive white noise current inputs; see also^53,55,58^. For each of the Control and GABA_B_R-AB models (see Results), we simulated the model for 50 trials of 12 s and used the last 10 s of simLFPs for the analysis^53,57^.

#### Pairwise neuronal correlations

To evaluate neuronal coordination within the model, we calculated the pairwise Spike Time Tiling Coefficient (STTC^59^) between all cell populations in the gamma-generating subnetwork (E-E, E-I, and I-I). The STTC was chosen as it provides a robust measure of spike synchrony that is independent of firing rate^59^, allowing for an unbiased comparison of network coordination across varying levels of excitatory drive and synaptic inhibition. To further ensure the significance of these neuronal correlations, we generated surrogate datasets using circular shuffling of spike trains (50 iterations per pair); for each cell pair, we subtracted the mean of its specific surrogate STTC distribution from the observed value. While the effect of this subtraction was negligible, it ensured that reported STTC values accounted for chance-level correlations. We utilized a synchrony time window of (Δt) of 5 ms, reflecting the precise temporal alignment characteristic of gamma-frequency rhythms. For each parameter set, STTC values were averaged across all possible pairs within the specified populations (E-E, E-I, and I-I) for each trial to generate the global pairwise synchrony maps (**Extended Data Fig. 5f**). Of note, since direct recurrent excitation between pyramidal cells is absent in the model (*G_EE_* = 0)—consistent with known CA1 anatomical circuitry—the observed E-E synchrony represents an emergent network property. This synchrony reflects the indirect entrainment of the excitatory population by the shared inhibitory feedback loop, wherein interneuronal activity provides the common temporal gating required to coordinate pyramidal cell spiking.

### Statistical Analysis

Statistical analyses were performed using R (v4.5.0), OriginPro 2019, GraphPad Prism 10, and MATLAB 2023a. Comprehensive statistical details are exhaustively reported in **Extended Data Table 2**, which includes specific tests utilized, exact sample sizes (n), exact p-values, and test statistics (e.g., *t*, *Z*, *F*, χ^2^, *U*) for each experiment. Unless otherwise stated, data are presented as mean ± SEM, and statistical significance was defined as *α* = 0.05.

Data normality was assessed using the Shapiro-Wilk test. For general comparisons between two independent groups, two-sample *t*-tests were applied to normally distributed data; otherwise, the non-parametric Mann-Whitney U-test was utilized. The permutation tests were performed based on the area under the curves with one million shuffles, as detailed in^53^. For experiments examining a single factor or two factors across multiple matched observations, one-way or two-way repeated measures ANOVAs were employed, respectively. For the comparison of FV vs. fEPSP amplitude, due to non-identical stimulation intensities, a linear regression model incorporating an interaction term (fEPSP ∼ FV * Group) was utilized to test for differences in regression slopes (interaction effect) and intercepts (main effect) between groups.

For multi-factorial designs involving repeated measures or nested data structures, Linear Mixed-Effects Models (LMMs) were employed using the *lme4* package. Experimental factors and their interactions were entered as fixed effects, while subject identity was modeled as a random intercept (e.g. 1 | Cell or 1 | Mouse) to properly account for within-subject correlations and tolerate unbalanced missing data. Omnibus testing of fixed effects was performed using Type III Wald chi-square tests via the *car* package, utilizing orthogonal contrasts to ensure accurate main effect estimations in the presence of interactions.

Where significant omnibus main effects or interactions were detected, comprehensive post-hoc simple main effects analyses were conducted using estimated marginal means (*emmeans* package). All resulting post-hoc pairwise comparisons were adjusted for multiple comparisons using the Benjamini-Hochberg (FDR) procedure to control the false discovery rate.

---

**Extended Data Fig. 1.**
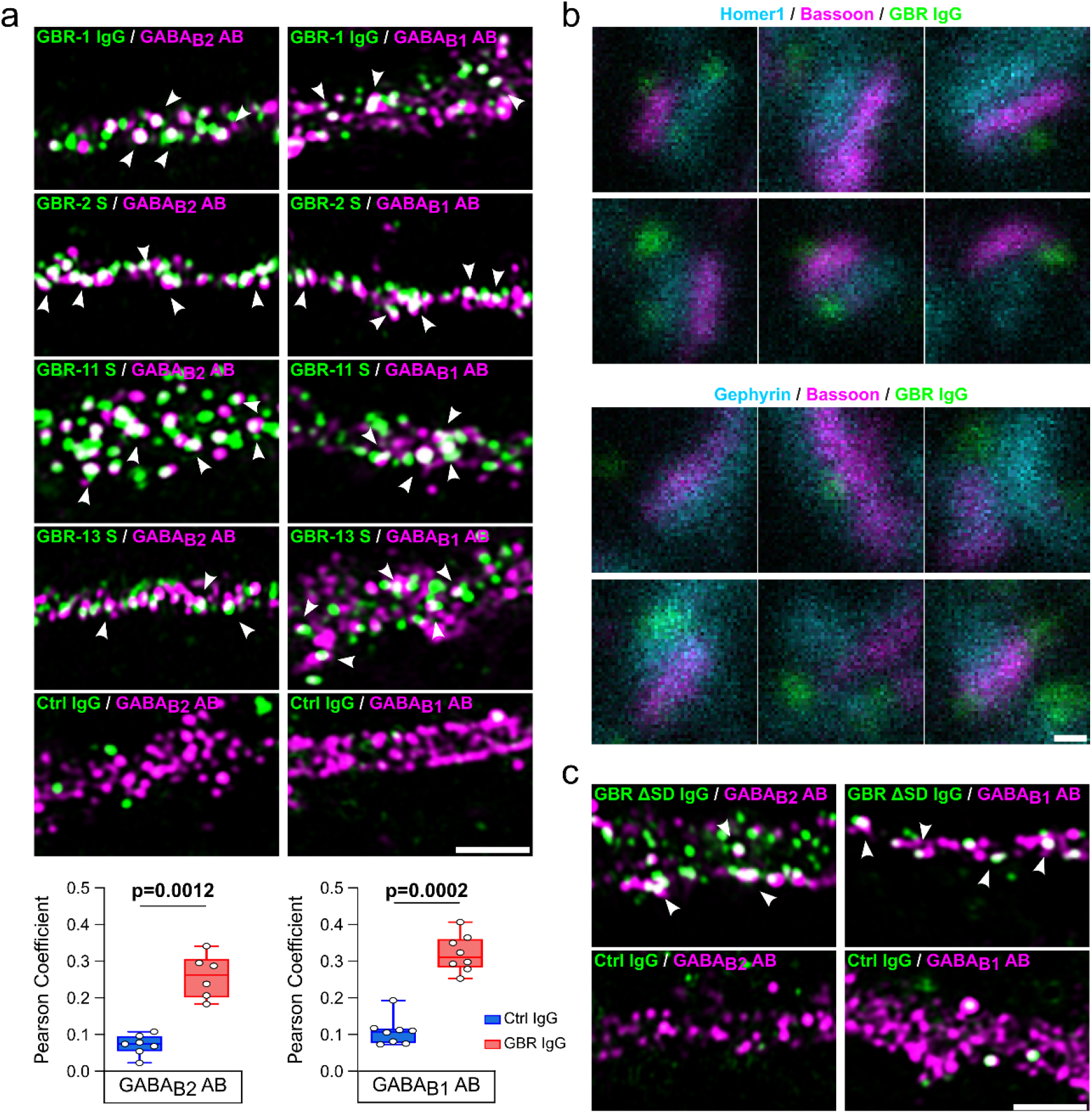
**a)** Upper rows: example confocal images of different anti-GABA_B_R purified IgG and serum samples with GABA_B_R antibodies (AB) to the GABA_B2_ and GABA_B1_ subunit (scale bar: 2 µm; colocalized puncta indicated by arrows). Bottom: colocalization quantified as Pearson’s Coefficient for purified GBR-1 IgG and Ctrl IgG (Ctrl IgG/GABA_B2_ AB n=7 vs. GBR IgG/GABA_B2_ AB n=8; Ctrl IgG/GABA_B1_ AB n=6 vs. GBR IgG/GABA_B1_ AB n=8; Mann-Whitney U-test). **b**) STED microscopy depicts perisynaptic localization of GBR IgG at glutamatergic synapses (Bassoon/Homer1; upper rows) and at GABAergic synapses (Bassoon/Gephyrin; lower rows). Scale bar: 100 nm. **c**) Example confocal images of GBR IgG depleted of the sushi-domain reactive portion (GBR-ΔSD IgG) and anti-GABA_B1_ or -GABA_B2_ ABs (scale bar: 2 µm; colocalized puncta indicated by arrows). Boxplots show median and interquartile range. Statistical details in **Extended Data Table 2**.

**Extended Data Fig. 2.**
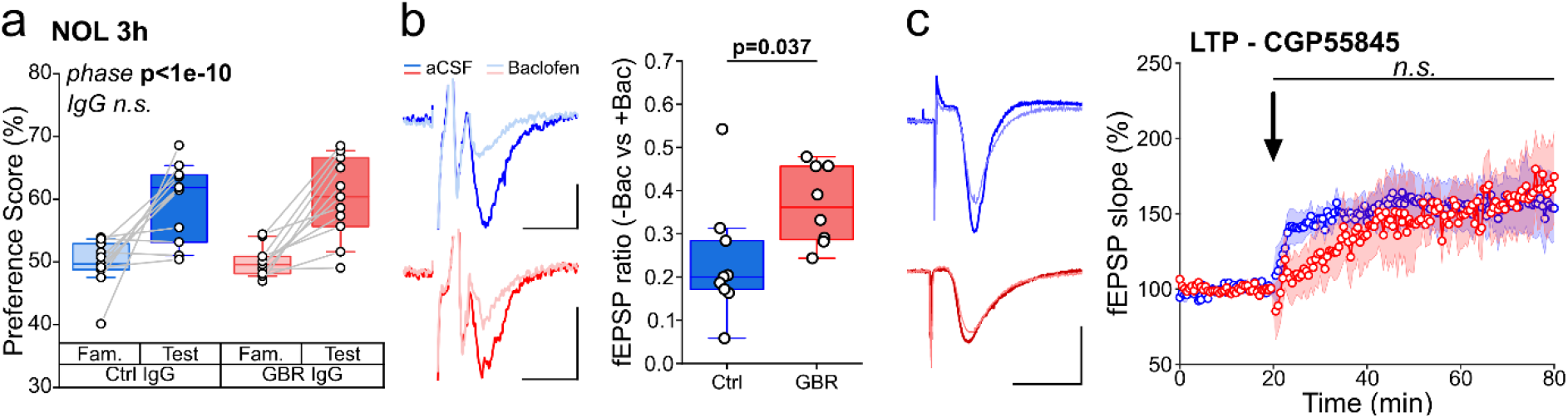
a) Ctrl IgG- and GBR IgG-treated mice show intact memory encoding in NOL test 3 h after familiarization phase (Fam. in graph; Ctrl IgG n=11, GBR IgG n=11; preference score ratios: Ctrl IgG 1.21 ± 0.06; GBR IgG 1.21 ± 0.04; LMM main effect analysis). **b)** Left: Example fEPSPs for Ctrl IgG (blue) and GBR IgG (red) before (dark shades) and after perfusion of baclofen (light shades; scale bars: 100 ms, 100 µV). Right: ratio of CA3 MF-induced fEPSP amplitudes before and after baclofen application showing a baclofen-induced reduction of fEPSP amplitude in Ctrl IgG-, but not in GBR IgG-treated mice (Ctrl IgG n=9, GBR IgG n=8; two-sample *t*-test). **c)** Left: example fEPSPs before (thin) and after TBS (bold; scale bars: 5 ms, 0.5 mV). Right: time course of fEPSP recording under CGP55845 shows a low LTP induction strength in Ctrl IgG- and GBR IgG-treated mice (Ctrl IgG n=8, GBR IgG n=7; whole-curve permutation test). Curves show mean ± SEM. Boxplots show median and interquartile range. *n.s.*, not significant. Statistical details in **Extended Data Table 2**.

**Extended Data Fig 3.**
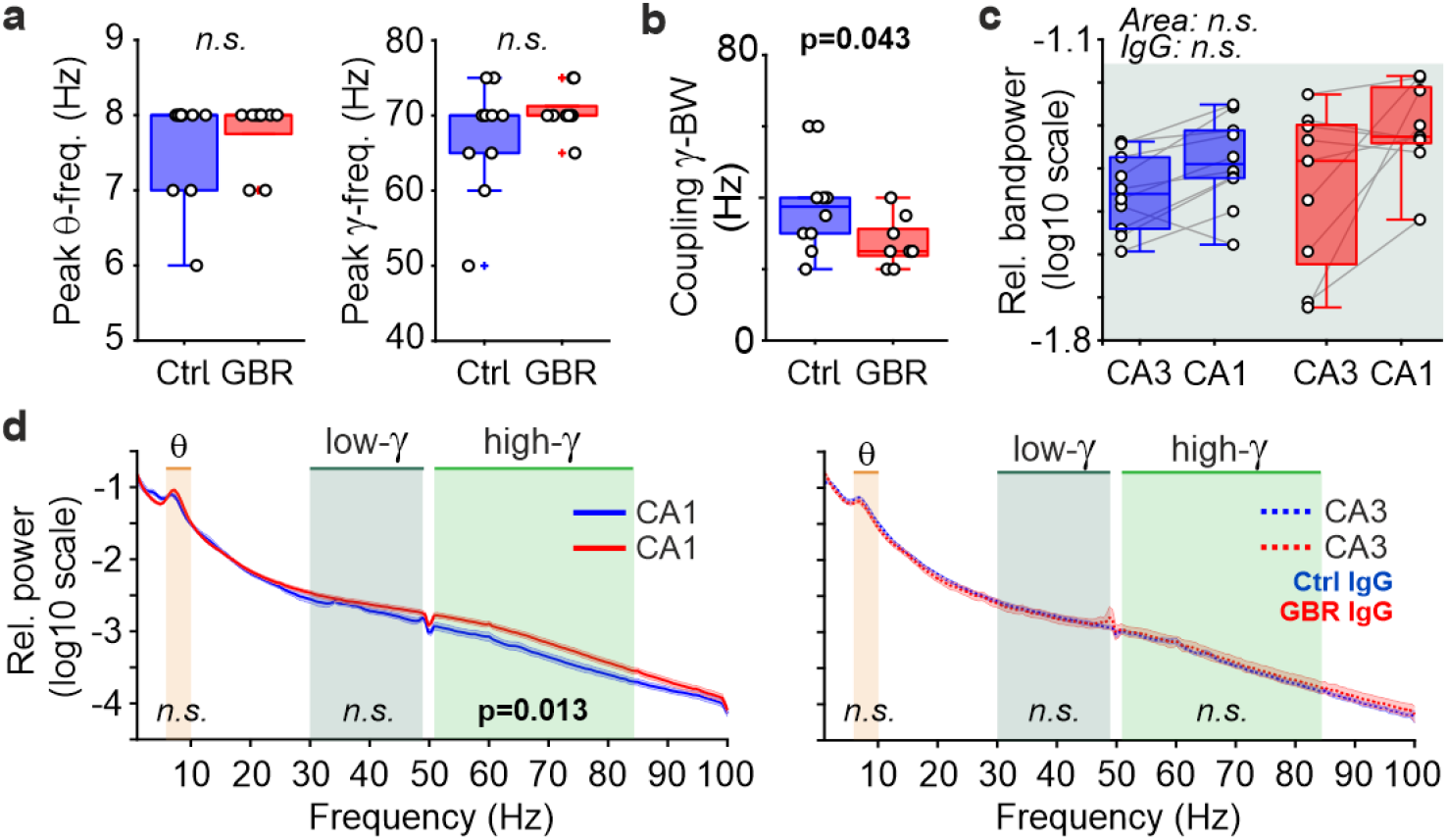
a) Peak theta and gamma frequencies of maximum coupling in the comodulogram. GBR IgG did not alter the peak coupling frequencies (Ctrl IgG n=9, GBR IgG n=10 mice; Mann-Whitney U-test). **b)** Coupling bandwidth in gamma frequency band (Ctrl IgG n=9, GBR IgG n=10 mice; Two-sample *t*-test) **c)** Relative bandpowers in the low-gamma band (30–50 Hz, right) remained unaltered by GBR IgG (Ctrl IgG n=9, GBR IgG n=10 mice; LMM main effect analysis). **d)** Mean relative power spectra, plotted as in Fig. 6d, but re-arranged to facilitate direct comparison between Ctrl IgG (blue) and GBR IgG (red) treatment groups within each hippocampal area. Left panel: CA1. Right: CA3. Note the increase in the high-gamma band (50–85 Hz) in CA1 under GBR IgG. The p-values for theta and high-gamma bands stem from statistics shown in Fig. 6e, and for low-gamma band from **(c)**. Curves show mean ± SEM. Boxplots show median and interquartile range. *n.s.*, not significant. Statistical details in **Extended Data Table 2**.

**Extended Data Fig 4.**
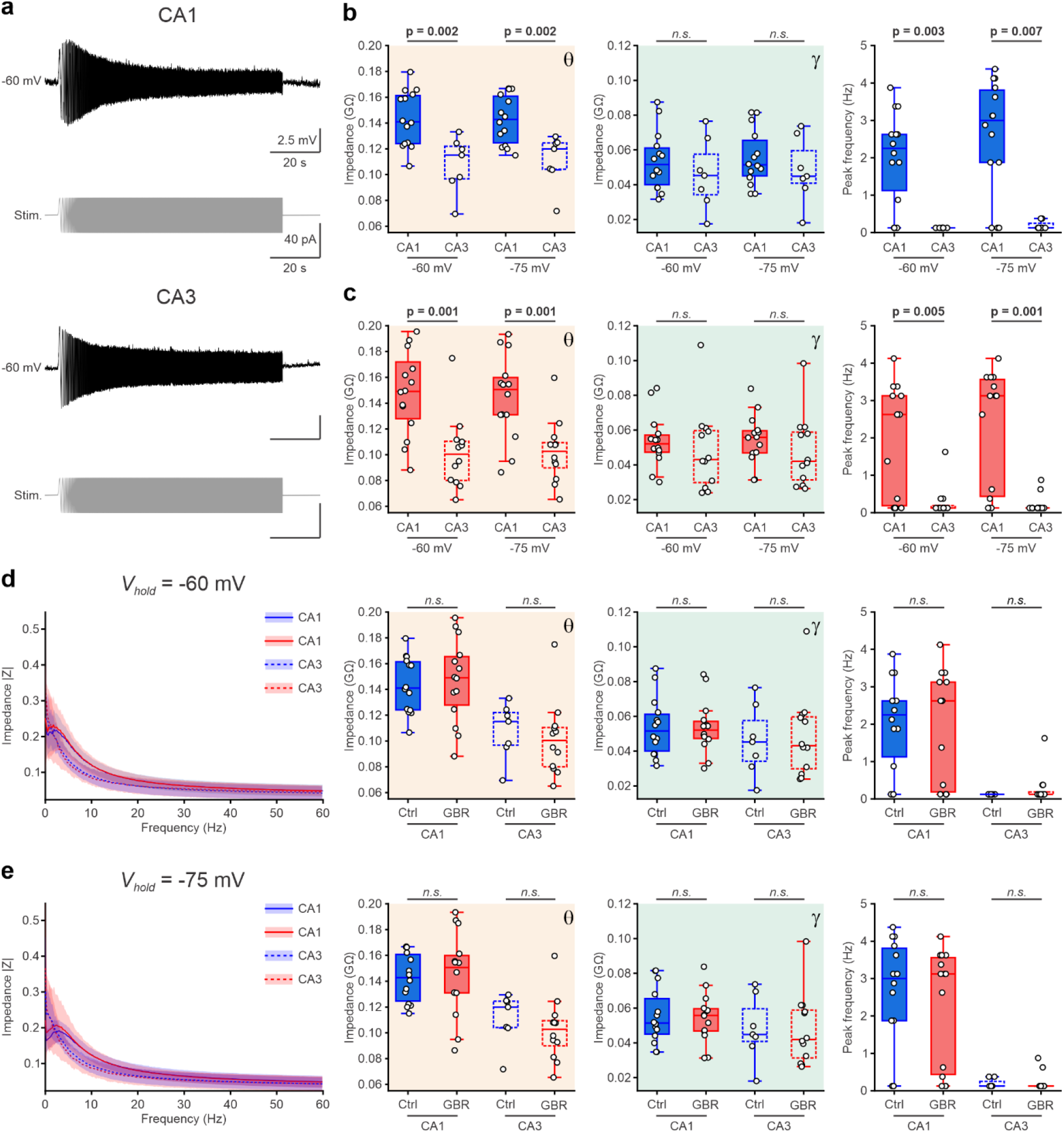
Subthreshold impedance profiles differ between CA1 and CA3 pyramidal cells and are not affected by GBR IgG. **a**) Representative ZAP recordings of CA1 (top) and CA3 (bottom) pyramidal cells (PCs; *I_hold_* = −60 mV) from a Ctrl IgG-treated mouse. Upper traces show membrane voltage responses to sinusoidal current injection with linearly increasing frequency (0-60 Hz, 90 s; lower traces; black). **b**) Subthreshold impedance comparison between CA1 and CA3 PCs in Ctrl IgG-treated mice (blue) *ex vivo*. Left: Impedance magnitudes (|Z|) in theta range (4–12 Hz, orange background). Middle: Impedance magnitudes in gamma range (30–60 Hz, dark green background). Right: Frequency preference of CA1 and CA3 PCs. CA1 and CA3 are shown as solid and dashed box plots, respectively. Each cell of was measured at two holding potentials of −60 mV and −75 mV. **c**) Same analysis as in (B) for GBR IgG-treated mice (red). **d**) Direct comparison of Ctrl IgG-(blue) and GBR IgG-treated mice (red) at *V_hold_* = −60 mV *ex vivo*. Left: Impedance profiles of CA1 (solid lines) and CA3 (dashed lines). Right: impedance magnitudes in theta (orange background) and gamma (dark green background) frequency bands, and peak (resonance) frequency of CA1 and CA3 PCs. CA1 and CA3 are shown as solid and dashed box plots, respectively. **e**) Same analysis as in (d) for recordings at *V_hold_* = −75 mV. **(b-e)** Open symbols represent individual cells (Ctrl IgG: CA1 n=14, CA3 n=7; GBR IgG: CA1 n=14, CA3 n=12). Curves show mean ± s.d. Box plots show median and interquartile range. *n.s.*, not significant. Statistical comparisons: two-sample *t*-tests or Mann-Whitney U-tests, as appropriate. Statistical details in **Extended Data Table 2**.

**Extended Data Fig 5.**
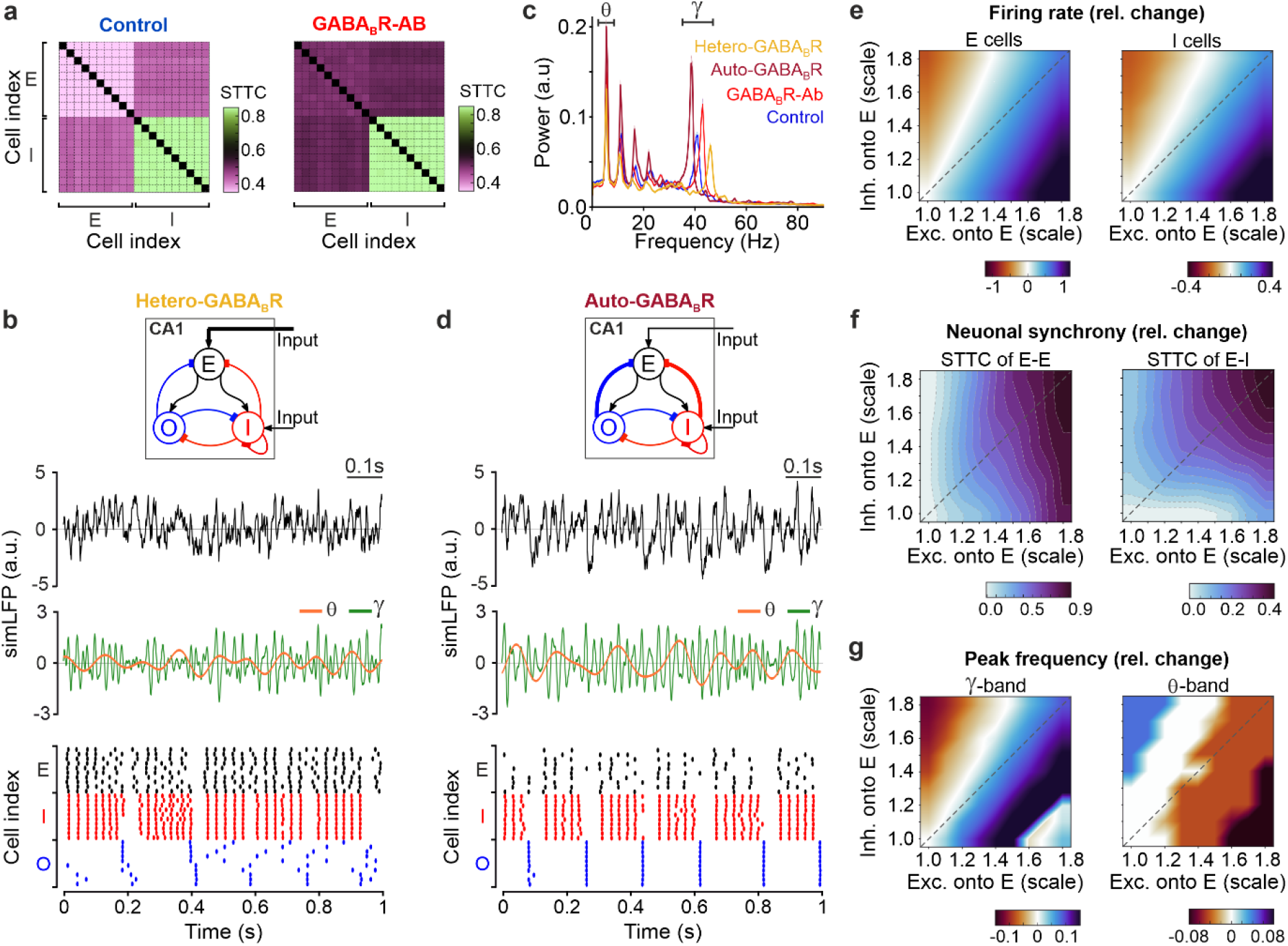
Distinct roles of auto- and hetero-GABA_B_R dysfunctions in amplified CA1 gamma oscillations. **a**) Pairwise Spike Time Tiling Coefficient (STTC) matrices for the Control and GABA_B_R-AB models shown in Fig. 7a**-b**. Darker colors indicate stronger neuronal correlation. The GABA_B_R-AB model exhibits increased synchrony within the E-cell population (top-left quadrant) and, crucially, between E- and I-cell populations (cross-diagonal quadrants), indicating augmented E-I coupling; thus, stronger gamma power (recall that the E-I subnetwork is the main generator of the simulated gamma rhythm). **b**) Hetero-GABA_B_R dysfunction model: Solely increasing excitatory drive onto E-cells (*ε_E_*; by 30%) leads to a state of excessive excitation. Spike rasters show elevated firing rates, but E-cells are only loosely time-locked to inhibitory feedback. This lack of precise temporal gating prevents the formation of coherent, high-amplitude gamma cycles, resulting in reduced network gamma power as compared to the Control model. **c**) Power spectra comparison. The Hetero-GABA_B_R model (yellow) shows a reduction in gamma power compared to the Control model, despite high excitation. The Auto-GABA_B_R model (brown) shows elevated theta and gamma power but a shift toward slower gamma frequencies. This reflects a state of inhibitory dominance resulting from increased O→E and I→E synaptic weights, which contributes to the amplified theta and gamma rhythms, respectively. Only the combined GABA_B_R-AB model (red) reproduces the high-frequency, high-power gamma phenotype observed *in vivo*. **d**) Auto-GABA_B_R dysfunction model: Solely increasing inhibitory synaptic weights onto E-cells (*G_OE_* and *G_IE_*; by 30%) creates a state of augmented inhibitory recruitment. Simulated LFPs (simLFPs) show strong oscillations, while spike rasters reveal sparse firing and a slower population gamma rhythm as compared to the Control model. **e-g)** Parameter space maps of network mechanisms. Values represent the relative change in parameter values compared to the Control model (coordinate 1.0, 1.0; same format as in Fig. 7d). **e**) Mean firing rates of E- and I-cells. Note that the GABA_B_ heteroreceptor dysfunction regime (lower diagonal) exhibits the highest firing rates but the lowest gamma power (cf. Fig 7d, top), demonstrating that elevated firing rates do not necessarily lead to gamma rhythm hypersynchrony. **f**) Neuronal synchrony (STTC). E-I synchrony is primarily governed by inhibition strength (vertical gradient), indicating that feedback inhibition regulates the entrainment of the pyramidal-interneuronal gamma (PING) loop. E-E synchrony increases with both E-drive and synaptic inhibition (diagonal gradient), reflecting increased population recruitment and rhythmic entrainment, respectively. Note the higher–more than double–range of the relative changes in E-E (0–0.9) as compared to E-I (0–0.4), indicating that excitatory synchronization (E-E) scales disproportionately thus failing to be effectively entrained by inhibitory feedback. Consequently, excessive E-E synchrony alone is insufficient to generate high gamma power (as seen in the hetero-GABA_B_R model); it requires a synergistic, parallel increase in E-I coupling to properly organize this excessive excitation into coherent gamma oscillations. **g**) Peak oscillation frequency. Gamma frequency scales with E-drive (horizontal gradient), indicating that E-drive acts as a frequency modulator, while inhibition provides the necessary temporal pacing. In contrast, theta frequency remains mostly stable across the parameter space (right panel), reflecting the robust pacing provided by the intrinsic properties of O-LM cells.

