## Extended Data Table 1 containing patient characteristics. for "Anti-GABA_B_ receptor autoimmunity causes cognitive decline by presynaptic pathology"

Patient characteristics

| Patient sample ID | Clinical Symptoms | Disease duration | $GABA_BR$ AB titer in serum (CBA) | $GABA_BR$ AB titer in CSF (CBA) | Coexisting autoantibodies | CSF lymphocytes n/µl | CSF protein mg/l | Concomitant tumor | Therapy |
|---|---|---|---|---|---|---|---|---|---|
| GBR IgG 1 | Limbic encephalitis with focal and generalized seizures; status epilepticus anterograde memory dysfunction; autonomic dysregulation | 1 month | 1:1000 | 1:320 | none | 30 | 587 | SCLC | Methylprednisolone, PLEX, RTX, IVIG |
| GBR S2 | Disorientation; memory impairment; psychosis with nocturnal aggressiveness; bilateral tonic-clonic seizures | 18 months | 1:320 | 1:32 | none | 1 | 458 | SCLC | Methylprednisolone, PLEX, RTX |
| GBR S3 | Epileptic seizures; cognitive impairment | 6 months | 1:10.000 | 1:320 | none | 43 | 430 | SCLC | Methylprednisolone |
| GBR S4 | Limbic encephalitis; status epilepticus; bulbar symptoms; memory dysfunction | 2 months | 1:100 | 1:100 | anti-SOX1, anti-Hu, anti-Zic4 | 29 | 410 | SCLC | Methylprednisolone, IVIG |
| GBR S5 | Severe gait disorder; sensory atactic neuropathy; tactile myoclonus; cognitive impairment | 6 days | 1:320 | 1:32 | anti-amphiphysin | 23 | 1705 | SCLC | Chemotherapy (Carboplatin, Etoposide, Durvalumab) for SCLC, no immunotherapy |
| GBR S6 | Limbic encephalitis with focal and generalized seizures, anterograde memory dysfunction, sleep disturbances, hyponatremia | 6 weeks | 1:100 | Not documen-ted | none | Not documented | Not documented | none | Methylprednisolone, IVIG, Mycofenolate-Mofetil |
| GBR S7 | Epileptic seizures; myoclonus; cognitive impairment | 8 months | 1:100 | 1:10 | none | 7 | 780 | LCNEC | Methylprednisolone |

| | | | | | | | | | |
|---|---|---|---|---|---|---|---|---|---|
| GBR S8 | Limbic encephalitis with focal temporal lobe seizures and memory disturbance; neuropathy; optic neuritis | 5 months | 1:10 | negative | anti-Zic4<br>anti-CV2<br>anti-SOX1<br>anti-Hu | 17 | 675 | SCLC | Chemotherapy for SCLC, no immunotherapy |
| GBR S9 | Limbic encephalitis with anterograde memory deficits and cognitive dysfunction; temporal lobe and secondary generalized seizures | 10 days | 1:100 | 1:32 | none | 71 | 694 | none | Methylprednisolone, IA |
| GBR S10 | Focal and complex partial epileptic seizures | 18 months | 1:10 | 1:32 | none | 135 | 646 | none | Methylprednisolone |
| GBR S11 | Limbic encephalitis with focal temporal lobe and generalized seizures; memory disturbance | 4 months | 1:10 | negative | none | 0 | 751 | none | Methylprednisolone, MTX |
| GBR S12 | Limbic encephalitis with anterograde memory dysfunction and deficits in executive functions; depression; weight loss; sleep abnormalities | 14 months | 1:100 | 1:10 | anti-IgLON5 | 2 | 617 | none | Methylprednisolone, PLEX, Cyclophosphamide |
| GBR S13 | Limbic encephalitis with anterograde memory dysfunction; double vision; gait disturbance | 2 months | 1:100 | negative | none | 0 | 614 | none | Methylprednisolone |
| GBR S14 | autoimmune encephalitis with seizures and retrograde amnesia, fully recovered; relapse with headache and mild memory deficit | 6 years; relapse 2 weeks | 1:100 | 1:1 | anti-NMDAR 1:10 in serum (CSF negative) | 263 | 521 | none | Methylprednisolone |
| GBR S15 | Brain stem syndrome, dysarthria, dysphagia, ataxia, gait disturbance, respiratory insufficiency | 3 days | 1:100 | Not dcoumen-ted | none | 3 | 610 | Oesophageal carcinoma (adeno carcinoma) | Radiotherapy; methylprednisolone, PLEX |

**Abbreviations:** Anti-Hu – antineuronal nuclear antibody-type 1 (ANNA-1); CSF – cerebrospinal fluid; CV2 – Collapsin Response Mediator Protein 5 (CRMP5); GBR – $GABA_BR$; IA – immunoadsorption; IgG – immunoglobulin G; IgLON5 – immunoglobulin-like cell adhesion molecule 5; IVIG – intravenous immunoglobulin; LCNEC – neuroendocrine lung carcinoma;

MTX – Methotrexat; NMDAR – N-methyl-D-aspartate (NMDA) receptor; PLEX – plasma exchange; RTX – Rituximab; S – serum; SCLC – small cell lung cancer; SOX1 – Sry-like high mobility group 1 (also: AGNA – Anti-Glial Nuclear Antibody); Zic4 – zinc finger protein 4
