## Extended Data Table 2 containing synopsis of statistical tests. for "Anti-GABA_B_ receptor autoimmunity causes cognitive decline by presynaptic pathology"

| Related to | Descriptive statistics | Test(s) | Test statistics |
|---|---|---|---|
| **Figure 1** | **Group: mean ± SEM (n = no. of cells; N = no. of mice)** | | |
| **1c** | Ctrl IgG VGlut: 0.01748 ± 0.005606 (n=12) | Linear Mixed-Effects Model (LMM) | |
| Colocalization Ratio | GBR IgG VGlut: 0.1716 ± 0.01985 (n=14) | Main effect of IgG: Ctrl vs. GBR | P = 1.637e-09 |
| | Ctrl IgG Homer1: 0.07043 ± 0.01058 (n=12) | (Type III Wald $\chi 2$) | $\chi^2_{(1)}$ = 36.3645 |
| | GBR IgG Homer1: 0.3231 ± 0.05473 (n=14) | Main effect of Marker: Homer1 vs. VGlut | P = 0.21113 |
| | | (Type III Wald $\chi 2$) | $\chi^2_{(1)}$ = 1.5637 |
| | | Interaction ([IgG] x [Marker]) | P = 0.08779 |
| | | (Type III Wald $\chi 2$) | $\chi^2_{(1)}$ = 2.9145 |
| **1d** | Ctrl IgG VGAT: 0.01707 ± 0.007621 (n=7) | Linear Mixed-Effects Model (LMM) | |
| Colocalization Ratio | GBR IgG VGAT: 0.1686 ± 0.02801 (n=10) | Main effect of IgG: Ctrl vs, GBR | P = 1.508e-06 |
| | Ctrl IgG Gephyrin: 0.09675 ± 0.04754 (n=6) | (Type III Wald $\chi 2$) | $\chi^2_{(1)}$ = 23.1378 |
| | GBR IgG Gephyrin: 0.3574 ± 0.04454 (n=10) | Main effect of Marker: Gephyrin vs. VGAT | P = 0.17232 |
| | | (Type III Wald $\chi 2$) | $\chi^2_{(1)}$ = 1.8626 |
| | | Interaction ([IgG] x [Marker]) | P = 0.14516 |
| | | (Type III Wald $\chi 2$) | $\chi^2_{(1)}$ = 2.1223 |
| **1e (top)** | Ctrl IgG aCSF: 3.4864 ± 0.7502 (n=9) | Mann-Whitney U-test (exact), two-tailed | P=0.003996 |
| Frequency (Hz) | Ctrl IgG Baclofen: 1.192 ± 0.1617 (n=5) | | Z=2.66667 |
| | | | U=43 |
| **1e (bottom)** | GBR IgG aCSF: 3.350 ± 0.8563 (n=7) | Mann-Whitney U-test (exact), two-tailed | P=0.56061 |
| Frequency (Hz) | GBR IgG Baclofen: 2.4778 ± 0.5378 (n=5) | | Z=0.56939 |
| | | | U=21.5 |
| **1f (bottom-left)** | Ctrl IgG: 1.155 ± 0.2817 (n=15) | Kruskal-Wallis ANOVA | P<0.0001 |
| Volume ($\mu m^3$) | 003-102 IgG: 7.811 ± 1.309 (n=18) | | $\chi^2$=36.51 |
| | GBR IgG: 9.540 ± 1.366 (n=16) | | df=2 |
| | Ctrl IgG vs 003-102 IgG | Dunn's post-hoc correction | Adj. P<0.0001 |
| | | | Z=4.947 |
| | Ctrl IgG vs GBR IgG | Dunn's post-hoc correction | Adj. P<0.0001 |
| | | | Z=5.526 |
| | 003-102 IgG vs GBR IgG | Dunn's post-hoc correction | Adj. P>0.9999 |
| | | | Z=0.7468 |
| **1f (bottom-right)** | 003-102 IgG: 1.351 ± 0.1753 (n=18) | Mann-Whitney U-test (exact), two-tailed | P<0.0001 |
| Ratio (Volume extracellular/intracellular) | GBR IgG: 6.444 ± 1.003 (n=16) | | Z=-4.76 |
| | | | U=6 |
| **Extended data Figure 1** | **Group: mean ± SEM (n = no. of cells; N = no. of mice)** | | |
| **1a (bottom-left)** | Ctrl IgG GABAB2 AB: 0.07175 ± 0.01052 (n=7) | Mann-Whitney U-test (exact), two-tailed | P=0.0012 |
| Pearson Coefficient | GBR IgG GABAB2 AB: 0.2585 ± 0.02435 (n=6) | | Z=-3.0 |
| | | | U=0 |
| **1a (bototm-right)** | Ctrl IgG GABAB1 AB: 0.1085 ± 0.01362 (n=8) | Mann-Whitney U-test (exact), two-tailed | P=0.0002 |
| Pearson Coefficient | GBR IgG GABAB1 AB: 0.3207 ± 0.01784 (n=8) | | Z=-3.36 |
| | | | U=0 |
| **Figure 2** | **Group: mean ± SEM (n = no. of cells; N = no. of mice)** | | |
| **2b** | Ctrl IgG Familiarization: 51.62 ± 1.647 (N=7) | Linear Mixed-Effects Model (LMM) | |
| Preference score (NOR 3h) | GBR IgG Familiarization: 50.31 ± 1.299 (N=8) | Main effect of IgG: Ctrl vs. GBR | P = 0.4389 |
| | Ctrl IgG Test: 59.21 ± 4.474 (N=7) | (Type III Wald $\chi 2$) | $\chi^2_{(1)}$ = 0.5990 |
| | GBR IgG Test: 64.88 ± 2.618 (N=8) | Main effect of Phase: Familirization vs. Test | P = 2.123e-05 |
| | | (Type III Wald $\chi 2$) | $\chi^2_{(1)}$ = 18.0759 |
| | | Interaction ([IgG] x [Phase]) | P = 0.1801 |
| | | (Type III Wald $\chi 2$) | $\chi^2_{(1)}$ =1.7966 |
| **2c** | Ctrl IgG Familiarization: 50.67 ± 0.4257 (N=10) | Linear Mixed-Effects Model (LMM) | |
| Preference score (NOR 24h) | GBR IgG Familiarization: 48.73 ± 0.7116 (N=9) | Main effect of IgG: Ctrl vs. GBR | P = 0.0002 |
| | Ctrl IgG Test: 66.46 ± 2.256 (N=10) | (Type III Wald $\chi 2$) | $\chi^2_{(1)}$ = 13.8276 |
| | GBR IgG Test: 55,44 ± 2.542 (N=9) | Main effect of Phase: Familirization vs. Test | P = 1.023e-10 |
| | | (Type III Wald $\chi 2$) | $\chi^2_{(1)}$ = 41.7774 |
| | | Interaction ([IgG] x [Phase]) | P = 0.0090 |
| | | (Type III Wald $\chi 2$) | $\chi^2_{(1)}$ = 6.8193 |
| | | Post-hoc simple effects (FDR corrected) | |
| | | Ctrl vs. GBR (Familiarization) | P = 0.4377 |
| | | LMM contrast (*t*-test) | $t_{(34)}$ = 0.785 |
| | | Ctrl vs. GBR (Test) | P = 0.0002 |
| | | LMM contrast (*t*-test) | $t_{(34)}$ = 4.476 |
| | | Familiarization vs. Test (Ctrl Group) | P = <0.0001 |
| | | LMM contrast (*t*-test) | $t_{(17)}$ = -6.593 |
| | | Familiarization vs. Test (GBR Group) | P = 0.0222 |
| | | LMM contrast (*t*-test) | $t_{(17)}$ = -2.655 |
| **Figure 3** | **Group: mean ± SEM (n = no. of cells; N = no. of mice)** | | |
| **3b (left)** | aCSF: (n=20, N=9) | two-way repeated ANOVA | P=9.69169E-5 |
| IPCS amplitude (normalized) Ctrl IgG | CGP (n=20; N=9) | | F=24.12219 |
| | | | df=1 |
| **3b (right)** | aCSF: (n=17, N=8) | two-way repeated ANOVA | P=0.33637 |
| IPCS amplitude (normalized) GBR IgG | CGP (n=17; N=8) | | F=0.98235 |
| | | | df=1 |
| **3d** | aCSF: (n=15, N=5) | two-way repeated ANOVA | P=3.27719E-5 |
| IPCS amplitude (normalized) GBR-ΔSD IgG | CGP (n=15; N=5) | | F=35.952 |
| | | | df=1 |
| **3f** | Ctrl IgG: (n=6; N=4) | one-way repeated ANOVA | P=0.00996 |
| EPSC Amplitude (pA) | aCSF: -184,0499 ± 36,34775 | | F=7.56858 |
| | Baclofen: -84,81801 ± 19,52409 | | df=2 |
| | CGP: -201,89169 ± 48,21797 | | |
| | aCSF vs Baclofen | Tukey post-hoc pairwise comparison | P=0.02962 |
| | | | t=4.32806 |
| | | | df=10 |
| | aCSF vs CGP | Tukey post-hoc pairwise comparison | P=0.84867 |
| | | | t=0.77818 |
| | | | df=10 |
| | Bclofen vs CGP | Tukey post-hoc pairwise comparison | P=0.01206 |
| | | | t=5.10625 |
| | | | df=10 |
| | GBR IgG: (n=9; N=4) | one-way repeated ANOVA | P=0.08909 |
| | aCSF: -138,0629 ± 18,3489 | | F=2.82331 |
| | Baclofen: -107,6420 ± 18,9150 | | df=2 |
| | CGP: -115,8183 ± 19,1763 | | |
| | aCSF vs Baclofen | Tukey post-hoc pairwise comparison | P=0.08539 |
| | | | t=3.2468 |
| | | | df=16 |

| | | | |
|---|---|---|---|
| | aCSF vs CGP | Tukey post-hoc pairwise comparison | P=0.24342<br>t=2.37415<br>df=16 |
| | Bclofen vs CGP | Tukey post-hoc pairwise comparison | P=0.81299<br>t=0.87265<br>df=16 |
| **3g**<br>Paired-pulse ratio | Ctrl IgG: (n=6; N=4)<br>aCSF: 1,44359 ± 0,06486<br>Baclofen: 1,72486 ± 0,09862<br>CGP: 1,37104 ± 0,02152 | one-way repeated ANOVA | P=0.02246<br>F=5.68298<br>df=2 |
| | aCSF vs Baclofen | Tukey post-hoc pairwise comparison | P=0.06975<br>t=3.58773<br>df=10 |
| | aCSF vs CGP | Tukey post-hoc pairwise comparison | P=0.79414<br>t=0.92553<br>df=10 |
| | Baclofen vs CGP | Tukey post-hoc pairwise comparison | P=0.02389<br>t=4.51326<br>df=10 |
| | GBR IgG: (n=9; N=4)<br>aCSF: 1,53548 ± 0,09387<br>Baclofen: 1,56342 ± 0,08664<br>CGP: 1,43849 ± 0,08467 | one-way repeated ANOVA | P=0.21009<br>F=1.72275<br>df=2 |
| | aCSF vs Baclofen | Tukey post-hoc pairwise comparison | P=0.91781<br>t=0.55939<br>df=16 |
| | aCSF vs CGP | Tukey post-hoc pairwise comparison | P=0.37781<br>t=1.94146<br>df=16 |
| | Baclofen vs CGP | Tukey post-hoc pairwise comparison | P=0.21162<br>t=2.50086<br>df=16 |
| **Figure 4** | **Group: mean ± SEM (n = no. of cells; N = no. of mice)** | | |
| **4b (left)**<br>R input (MΩ) | Ctrl IgG: 181,80758 ± 12,27863 (n=13; N=3)<br>GBR IgG: 162,78831 ± 14,20228 (n=11; N=3) | two-sample t-test | P=0.31949<br>t=1.01854<br>df=22 |
| **4b (right)**<br>membrane tau τ (ms) | Ctrl IgG: 24,46255 ± 1,7195 (n=13; N=3)<br>GBR IgG: 25,63982 ± 1,97206 (n=11; N=3) | Mann-Whitney U-test (exact), two-tailed | P= 0.73296<br>Z=-0.34762<br>U=65 |
| **4e**<br>sIPSP amplitude (mV) | Ctrl IgG: -4,9324 ± 0,55305 (n=18; N=5)<br>GBR IgG: -4,76532 ± 0,44081 (n=12; N=5) | Mann-Whitney U-test (exact), two-tailed | P= 0.81874<br>Z=0.23283<br>U=114 |
| **4f (middle)**<br>Latency to 1st AP (ms) | Ctrl IgG -HFS: 105,65467 ± 7,28992 (n=18; N=5)<br>Ctrl IgG +HFS: 128,64957 ± 7,68893 (n=18; N=5)<br>GBR IgG -HFS: 96,417 ± 10,77742 (n=12; N=5)<br>GBR IgG +HFS: 116,71081 ± 8,78871 (n=12; N=5) | Linear Mixed-Effects Model (LMM) | |
| | | Main effect of IgG: Ctrl vs. GBR<br>(Type III Wald χ2) | P = 0.3723<br>$\chi^2_{(1)}$ = 3.3885 |
| | | Main effect of HFS: -HFS vs. +HFS<br>(Type III Wald χ2) | P = 5.155e-15<br>$\chi^2_{(1)}$ = 61.2005 |
| | | Interaction ([IgG] x [HFS])<br>(Type III Wald χ2) | P = 0.6255<br>$\chi^2_{(1)}$ = 0.2383 |
| **4f (right)**<br>Number of APs | Ctrl IgG -HFS: 4,43889 ± 0,35074 (n=18; N=5)<br>Ctrl IgG +HFS: 3,30873 ± 0,31885 (n=18; N=5)<br>GBR IgG -HFS: 5,49167 ± 0,57754 (n=12; N=5)<br>GBR IgG +HFS: 4,32593 ± 0,41137 (n=12; N=5) | Linear Mixed-Effects Model | |
| | | Main effect of IgG: Ctrl vs. GBR<br>(Type III Wald χ2) | P = 0.06565<br>$\chi^2_{(1)}$ = 3.3885 |
| | | Main effect of HFS: -HFS vs. +HFS<br>(Type III Wald χ2) | P < 2e-16<br>$\chi^2_{(1)}$ = 69.3894 |
| | | Interaction ([IgG] x [HFS])<br>(Type III Wald χ2) | P = 0.89728<br>$\chi^2_{(1)}$ = 0.0167 |
| **4h**<br>Spiking probability | Ctrl IgG:<br>aCSF -HFS: 0,8625 ± 0,03441 (n=10; N=4)<br>aCSF +HFS: 0,4975 ± 0,08014 (n=10; N=4)<br>CGP -HFS: 0,94167 ± 0,04167 (n=6; N=4)<br>CGP +HFS: 0,8625 ± 0,06511 (n=6; N=4)<br><br>GBR IgG:<br>aCSF -HFS: 0,83929 ± 0,03562 (n=14; N=5)<br>aCSF +HFS: 0,525 ± 0,07927 (n=14; N=5)<br>CGP -HFS: 0,91875 ± 0,03399 (n=8; N=3)<br>CGP +HFS: 0,7625 ± 0,07778 (n=8; N=3) | 3-Way Linear Mixed-Effects Model (LMM) | |
| | | Main effect of IgG: Ctrl vs. GBR<br>(Type III Wald χ2) | P = 0.53173<br>$\chi^2_{(1)}$ = 0.3911 |
| | | Main effect of Solution: aCSF vs. CGP<br>(Type III Wald χ2) | P = 6.012e-05<br>$\chi^2_{(1)}$ = 16.0990 |
| | | Main effect of HFS: -HFS vs. +HFS<br>(Type III Wald χ2) | P = 1.422e-06<br>$\chi^2_{(1)}$ = 23.2509 |
| | | Interaction ([IgG] x [Solution])<br>(Type III Wald χ2) | P = 0.4496<br>$\chi^2_{(1)}$ = 0.50250 |
| | | Interaction ([IgG] x [HFS])<br>(Type III Wald χ2) | P = 0.88945<br>$\chi^2_{(1)}$ = 0.0193 |
| | | Interaction ([Solution] x [HFS])<br>(Type III Wald χ2) | P = 0.01929<br>$\chi^2_{(1)}$ = 5.4751 |
| | | Interaction ([IgG] x [Solution] x [HFS])<br>(Type III Wald χ2) | P = 0.50051<br>$\chi^2_{(1)}$ = 0.4539 |
| | | Post-hoc simple effects (FDR corrected) | |
| | | aCSF vs. CGP (-HFS, Ctrl IgG)<br>LMM contrast (*t*-test) | P = 0.4428<br>$t_{(55.1)}$ = -0.773 |
| | | aCSF vs. CGP (+HFS, Ctrl IgG)<br>LMM contrast (*t*-test) | P = 0.0030<br>$t_{(55.1)}$ = -3.564 |
| | | aCSF vs. CGP (-HFS, GBR IgG)<br>LMM contrast (*t*-test) | P = 0.4428<br>$t_{(55.8)}$ = -0.904 |
| | | aCSF vs. CGP (+HFS vs. GBR)<br>LMM contrast (*t*-test) | P = 0.0183<br>$t_{(55.8)}$ = -2.701 |
| | | -HFS vs. +HFS (aCSF, Ctrl IgG)<br>LMM contrast (*t*-test) | P = 0.0003<br>$t_{(48.1)}$ = 4.141 |
| | | -HFS vs. +HFS (CGP, Ctrl IgG)<br>LMM contrast (*t*-test) | P = 0.4900<br>$t_{(48.1)}$ = 0.696 |
| | | -HFS vs. +HFS (aCSF, GBR IgG)<br>LMM contrast (*t*-test) | P = 0.0003<br>$t_{(48.1)}$ = 4.219 |
| | | -HFS vs. +HFS (aCSF, GBR IgG)<br>LMM contrast (*t*-test) | P = 0.1592<br>$t_{(48.1)}$ = 1.585 |
| **Figure 5** | **Group: mean ± SEM (n = no. of cells; N = no. of mice)** | | |
| **5c**<br>Paired-pulse ratio | Ctrl IgG: 3,55822 ± 0,21814 (n=23; N=10)<br>GBR IgG: 2,92674 ± 0,14317 (n=25; N=10) | Mann-Whitney U-test (exact), two-tailed | P=0.02931<br>Z=2.16695<br>U=393 |
| **5d**<br>Input-Output curve (FV vs. fEPSP) | toral number of samples = 132 data points<br>Ctrl IgG: (n=23 slices; N=9)<br>GBR IgG (n=21 slices; N=8) | Linear Regression (Y ~ X * Group) | |
| | | Intercept difference (Main effect of IgG)<br>Regression coefficient (*t*-test) | P = 0.212<br>$t_{(128)}$ = -1.255 |
| | | Baseline slope (Main effect of FV in Ctrl) | P = 0.000112 |

| | | | |
|---|---|---|---|
| | | Regression coefficient ($t$-test) | $t_{(128)}$ = 3.987 |
| | | Slope difference (FV × IgG interaction)<br>Regression coefficient ($t$-test) | P = 0.0144<br>$t_{(128)}$ = 2.482 |
| **5f**<br>fEPSP slope (mV/ms) | Ctrl IgG: (n=31 slices; N=11)<br>GBR IgG (n=32 slices; N=11) | whole-curve permutation, two-tailed, 1 million shuffles | P=0.0092 |
| **5g**<br>fEPSP slope (%) | Ctrl IgG: (n=11 slices; N=8)<br>GBR IgG: (n=17 slices; N=11) | whole-curve permutation, two-tailed, 1 million shuffles | P=0.0096 |
| **Extended Data Figure 2** | **Group: mean ± SEM (n = no. of cells; N = no. of mice)** | | |
| **E2a**<br>Preference score (NOL 3h) | Ctrl IgG Familiarization: 49.76 ± 1.151 (N=11)<br>GBR IgG Familiarization: 50.04 ± 0.7277 (N=11)<br>Ctrl IgG Test: 59.70 ± 1.855 (N=11)<br>GBR IgG Test: 60.24 ± 1.960 (N=11) | Linear Mixed-Effects Model (LMM) | |
| | | Main effect of IgG: Ctrl vs. GBR<br>(Type III Wald $\chi$2) | P = 0.7853<br>$\chi^2_{(1)}$ = 0.0742 |
| | | Main effect of Phase: Familirization vs. Test<br>(Type III Wald $\chi$2) | P = 2.77e-11<br>$\chi^2_{(1)}$ = 44.3332 |
| | | Interaction ([IgG] x [Phase])<br>(Type III Wald $\chi$2) | P = 0.9322<br>$\chi^2_{(1)}$ = 0.0072 |
| **E2b**<br>fEPSP ratio (-Bac / +Bac) | Ctrl IgG: 0,23583 ± 0,04534 (n=9; N=5)<br>GBR IgG: 0,36626 ± 0,03232 (n=8; N=5) | two-sample t-test, two-tailed | P=0.03711<br>t=-2.28755<br>df=15 |
| **E2c**<br>fEPSP slope (%) | Ctrl IgG: (n=8 slices; N=7)<br>GBR IgG: (n=7 slices; N=6) | whole-curve permutation, two-tailed, 1 million shuffles | P= 0.6537 |
| **Figure 6** | **Group: mean ± SEM (n = no. of cells; N = no. of mice)** | | |
| **6c (left)**<br>Peak MI value | Ctrl IgG: 10.69145 ± 1.2218 (n=N/A; N=10)<br>GBR IgG: 15.46717 ± 1.61425 (n=N/A; N=9) | two-sample t-test, two-tailed | P = 0.0288<br>t = -2.38837<br>df = 17 |
| **6c (middle)**<br>Coupling area; the hotspot in comodulogram | Ctrl IgG: 32.6 ± 6.8754 (n=N/A; N=10)<br>GBR IgG: 17.77778 ± 1.63959 (n=N/A; N=9) | Mann-Whitney U-test (exact), two-tailed | P = 0.01057<br>Z = 2.45488<br>U = 75.5 |
| **6c (rigth)**<br>Bandwidth of coupling area in theta band | Ctrl IgG: 3.9 ± 0.45826 (n=N/A; N=10)<br>GBR IgG: 2.44444 ± 0.24216 (n=N/A; N=9) | Mann-Whitney U-test (exact), two-tailed | P = 0.01<br>Z = 2.68855<br>U = 75 |
| **6d (right)**<br>Overall power ratio (CA1 over CA3) | Ctrl IgG: 0.37418 ± 1.4282 (n=N/A; N=10) | one-Sample t-test, two-tailed | P = 0.79922<br>t = 0.26199<br>df = 9 |
| | GBR IgG: 4.08692 ± 1.57962 (n=N/A; N=9) | one-sample Wilcoxon signed-rank test (exact), two-tailed | P = 0.01953<br>Z = 2.25093<br>W = 42 |
| **6d (left)**<br>Relative bandpowers in specfic bands | (See statistics in 6e and E3c) | | |
| **6e (left)**<br>Relative bandpower in theta band | Ctrl IgG CA3: -0.5745 ± 0.0234 (n=N/A; N=10)<br>Gtrl IgG CA1: -0.5591 ± 0.0242 (n=N/A; N=10)<br>GBR IgG CA3: -0.6259 ± 0.0278 (n=N/A; N=9)<br>GBR IgG CA1: -0.5069 ± 0.0198 (n=N/A; N=9) | Generalized Linear Mixed-Effects Model (GLMM, Gamma distribution) | |
| | | Main effect of IgG: Ctrl vs. GBR<br>(Type III Wald $\chi$2) | P = 0.2064<br>$\chi^2_{(1)}$ = 1.5964 |
| | | Main effect of Area: CA1 vs. CA3<br>(Type III Wald $\chi$2) | P = 0.4675<br>$\chi^2_{(1)}$ = 0.5279 |
| | | Interaction ([IgG] x [Area])<br>(Type III Wald $\chi$2) | P = 0.0001087<br>$\chi^2_{(1)}$ = 14.98 |
| | | Post-hoc simple effects (FDR corrected) | |
| | | Ctrl vs. GBR (CA3)<br>GLMM contrast ($Z$-test) | P = 0.2752<br>$Z$ = 1.263 |
| | | Ctrl vs. GBR (CA1)<br>GLMM contrast ($Z$-test) | P = 0.2752<br>$Z$ = -1.438 |
| | | CA3 vs. CA1 (Ctrl IgG)<br>GLMM contrast ($Z$-test) | P = 0.4675<br>$Z$ = -0.727 |
| | | CA3 vs. CA1 (GBR IgG)<br>GLMM contrast ($Z$-test) | P < 0.0001<br>$Z$ = -5.953 |
| **6e (right)**<br>Relative bandpower in high-gamma band | Ctrl IgG CA3: -1.7628 ± 0.0232 (n=N/A; N=10)<br>Gtrl IgG CA1: -1.6801 ± 0.0515 (n=N/A; N=10)<br>GBR IgG CA3: -1.7336 ± 0.0752 (n=N/A; N=9)<br>GBR IgG CA1: -1.5055 ± 0.0405 (n=N/A; N=9) | Generalized Linear Mixed-Effects Model (GLMM, Gamma distribution) | |
| | | Main effect of IgG: Ctrl vs. GBR<br>(Type III Wald $\chi$2) | P = 0.8135<br>$\chi^2_{(1)}$ = 0.0557 |
| | | Main effect of Area: CA1 vs. CA3<br>(Type III Wald $\chi$2) | P = 0.1264<br>$\chi^2_{(1)}$ = 2.3363 |
| | | Interaction ([IgG] x [Area])<br>(Type III Wald $\chi$2) | P = 0.0005<br>$\chi^2_{(1)}$ = 12.1501 |
| | | Post-hoc simple effects (FDR corrected) | |
| | | Ctrl vs. GBR (CA3)<br>GLMM contrast ($Z$-test) | P = 0.8135<br>$Z$ = -0.236 |
| | | Ctrl vs. GBR (CA1)<br>GLMM contrast ($Z$-test) | P = 0.0129<br>$Z$ = -2.724 |
| | | CA3 vs. CA1 (Ctrl IgG)<br>GLMM contrast ($Z$-test) | P = 0.1685<br>$Z$ = -1.528 |
| | | CA3 vs. CA1 (GBR IgG)<br>GLMM contrast ($Z$-test) | P < 0.0001<br>$Z$ = -5.391 |
| **6f**<br>AUC of mean MI over theta-band curve at low-gamma (LG) and high-gamma (HG) bands | Ctrl IgG LG: 24.4003 ± 1.5795 (n=N/A; N=10)<br>Gtrl IgG HG: 52.2754 ± 5.8116 (n=N/A; N=10)<br>GBR IgG LG: 25.763 ± 1.2974 (n=N/A; N=9)<br>GBR IgG HG: 69.4611 ± 6.3202 (n=N/A; N=9) | Generalized Linear Mixed-Effects Model (GLMM, Gamma distribution) | |
| | | Main effect of IgG: Ctrl vs. GBR<br>(Type III Wald $\chi$2) | P = 0.56877<br>$\chi^2_{(1)}$ = 0.3247 |
| | | Main effect of Gamma Band: LG vs. HG<br>(Type III Wald $\chi$2) | P = 8.959e-12<br>$\chi^2_{(1)}$ = 46.5439 |
| | | Interaction ([IgG] x [Gamma Band])<br>(Type III Wald $\chi$2) | P = 0.01477<br>$\chi^2_{(1)}$ = 5.9434 |
| | | Post-hoc simple effects (FDR corrected) | |
| | | Ctrl vs. GBR (LG)<br>GLMM contrast ($Z$-test) | P = 0.5688<br>$Z$ = -0.570 |
| | | Ctrl vs. GBR (HG)<br>GLMM contrast ($Z$-test) | P = 0.0058<br>$Z$ = -2.851 |
| | | LG vs. HG (Ctrl IgG)<br>GLMM contrast ($Z$-test) | P < 0.0001<br>$Z$ = -6.822 |
| | | LG vs. HG (GBR IgG)<br>GLMM contrast ($Z$-test) | P < 0.0001<br>$Z$ = -8.049 |
| **6g**<br>Percentage of time bins with significant coupling | Gtrl IgG CA1: 46.05868 ± 7.83547 (n=N/A; N=10)<br>GBR IgG CA3: 46.40982 ± 7.76492 (n=N/A; N=9) | two-sample t-test, two-tailed | P = 0.97506<br>t = -0.03172<br>df = 17 |
| **Extended Data Figure 3** | **Group: mean ± SEM (n = no. of cells; N = no. of mice)** | | |
| **E3a (left)**<br>Theta-frequency at peak MI | Ctrl IgG: 7.6 ± 0.22111 (n=N/A; N=10)<br>GBR IgG: 7.77778 ± 0.14699 (n=N/A; N=9) | Mann-Whitney U-test (exact), two-tailed | P = 0.81424<br>Z = -0.42444<br>U = 40.5 |
| **E3a (right)**<br>Gamma-frequency at peak MI | Ctrl IgG: 67 ± 2.38048 (n=N/A; N=10)<br>GBR IgG: 70.55556 ± 1.00154 (n=N/A; N=9) | Mann-Whitney U-test (exact), two-tailed | P = 0.33443<br>Z = -1.02269 |

| | | | |
|---|---|---|---|
| | | | U = 33 |
| **E3b**<br>Bandwidth of coupling area in gamma band | Ctrl IgG: 38 ± 4.22953 (n=N/A; N=10)<br>GBR IgG: 27.222 ± 2.222 (n=N/A; N=9) | two-sample t-test, two-tailed | P = 0.04348<br>t = 2.1815<br>df = 17 |
| **E3c**<br>Relative bandpower in low-gamma band | Ctrl IgG CA3: -1.4569 ± 0.0284 (n=N/A; N=10)<br>Gtrl IgG CA1: -1.3842 ± 0.0328 (n=N/A; N=10)<br>GBR IgG CA3: -1.4484 ± 0.0622 (n=N/A; N=9)<br>GBR IgG CA1: -1.3074 ± 0.0347 (n=N/A; N=9) | Linear Mixed-Effects Model (LMM) | |
| | | Main effect of IgG: Ctrl vs. GBR<br>(Type III Wald $\chi 2$) | P = 0.57371<br>$\chi^2_{(1)}$ = 0.3165 |
| | | Main effect of Area: CA1 vs. CA3<br>(Type III Wald $\chi 2$) | P = 0.08497<br>$\chi^2_{(1)}$ = 2.9673 |
| | | Interaction ([IgG] x [Area])<br>(Type III Wald $\chi 2$) | P = 0.34845<br>$\chi^2_{(1)}$ = 0.8791 |
| **E3d**<br>Relative bandpowers in specfic bands | (See statistics in 6e and E3c) | | |
| **Extended Data Figure 4** | **Group: mean ± SEM (n = no. of cells; N = no. of mice)** | | |
| **E4b (left)**<br>Impedance \|Z\| (GΩ) | Ctrl IgG CA1 -60 mV: 0.1435 ± 0.0218 (n=14; N=3)<br>Ctrl IgG CA3 -60 mV: 0.1079 ± 0.0217 (n=7; N=3) | Two sample t-test, two-tailed | P = 0.0022<br>t = 3.5385<br>df = 19 |
| | Ctrl IgG CA1 -75 mV: 0.1426 ± 0.0190 (n=14; N=3)<br>Ctrl IgG CA3 -75 mV: 0.1112 ± 0.0201 (n=7; N=3) | Two sample t-test, two-tailed | P = 0.0024<br>t = 3.5034<br>df = 19 |
| **E4b (middle)**<br>Impedance \|Z\| (GΩ) | Ctrl IgG CA1 -60 mV: 0.0538 ± 0.0170 (n=14; N=3)<br>Ctrl IgG CA3 -60 mV: 0.0461 ± 0.0203 (n=7; N=3) | Two sample t-test, two-tailed | P = 0.3707<br>t = 0.9168<br>df = 19 |
| | Ctrl IgG CA1 -75 mV: 0.0554 ± 0.0161 (n=14; N=3)<br>Ctrl IgG CA3 -75 mV: 0.0482 ± 0.0189 (n=7; N=3) | Two sample t-test, two-tailed | P = 0.3737<br>t = 0.9109<br>df = 19 |
| **E4b (right)**<br>Peak frequency (Hz) | Ctrl IgG CA1 -60 mV: 2 ± 1.2548 (n=14; N=3)<br>Ctrl IgG CA3 -60 mV: 0.1250 ± 0 (n=7; N=3) | Mann-Whitney U-test, two-tailed | P = 0.0026<br>Z = 3.0068<br>U = 87.5 |
| | Ctrl IgG CA1 -75 mV: 2.5714 ± 1.5323 (n=14; N=3)<br>Ctrl IgG CA3 -75 mV: 0.1964 ± 0.1220 (n=7; N=3) | Mann-Whitney U-test, two-tailed | P = 0.0072<br>Z = 2.6891<br>U = 84.5 |
| **E4c (left)**<br>Impedance \|Z\| (GΩ) | GBR IgG CA1 -60 mV: 0.1467 ± 0.323 (n=14; N=5)<br>GBR IgG CA3 -60 mV: 0.1015 ± 0.0288 (n=12; N=3) | Two sample t-test, two-tailed | P = 0.0010<br>t = 3.7402<br>df = 24 |
| | GBR IgG CA1 -75 mV: 0.1448 ± 0.0325 (n=14; N=5)<br>GBR IgG CA3 -75 mV: 0.1025 ± 0.0245 (n=12; N=3) | Two sample t-test, two-tailed | P = 0.0011<br>t = 3.6899<br>df = 24 |
| **E4c (middle)**<br>Impedance \|Z\| (GΩ) | GBR IgG CA1 -60 mV: 0.0537 ± 0.0152 (n=14; N=5)<br>GBR IgG CA3 -60 mV: 0.0485 ± 0.0238 (n=12; N=3) | Mann-Whitney U-test, two-tailed | P = 0.3159<br>Z = 1.003<br>U = 104 |
| | GBR IgG CA1 -75 mV: 0.0546 ± 0.0144 (n=14; N=5)<br>GBR IgG CA3 -75 mV: 0.0479 ± 0.0208 (n=12; N=3) | Two sample t-test, two-tailed | P = 0.3442<br>t = 0.9649<br>df = 24 |
| **E4c (right)**<br>Peak frequency (Hz) | GBR IgG CA1 -60 mV: 1.9464 ± 1.4950 (n=14; N=5)<br>GBR IgG CA3 -60 mV: 0.2917 ± 0.4308 (n=12; N=3) | Mann-Whitney U-test, two-tailed | P = 0.0054<br>Z = 2.7813<br>U = 135 |
| | GBR IgG CA1 -75 mV: 2.2679 ± 1.5833 (n=14; N=5)<br>GBR IgG CA3 -75 mV: 0.2292 ± 0.2491 (n=12; N=3) | Mann-Whitney U-test, two-tailed | P = 0.0012<br>Z = 3.2488<br>U = 143,5 |
| **E4d (left)**<br>Impedance \|Z\| (GΩ) | Ctrl IgG CA1 -60 mV: 0.1435 ± 0.0218 (n=14; N=3)<br>GBR IgG CA1 -60 mV: 0.1467 ± 0.323 (n=14; N=5) | Two sample t-test, two-tailed | P = 0.7586<br>t = -0.3106<br>df = 26 |
| | Ctrl IgG CA3 -60 mV: 0.1079 ± 0.0217 (n=7; N=3)<br>GBR IgG CA3 -60 mV: 0.1015 ± 0.0288 (n=12; N=3) | Two sample t-test, two-tailed | P = 0.6188<br>t = 0.5069<br>df = 17 |
| **E4d (middle)**<br>Impedance \|Z\| (GΩ) | Ctrl IgG CA1 -60 mV: 0.0538 ± 0.0170 (n=14; N=3)<br>GBR IgG CA1 -60 mV: 0.0537 ± 0.0152 (n=14; N=5) | Two sample t-test, two-tailed | P = 0.9885<br>t = 0.0146<br>df = 26 |
| | Ctrl IgG CA3 -60 mV: 0.0461 ± 0.0203 (n=7; N=3)<br>GBR IgG CA3 -60 mV: 0.0485 ± 0.0238 (n=12; N=3) | Mann-Whitney U-test, two-tailed | P = 0.8991<br>Z = 0.1268<br>U = 44 |
| **E4d (right)**<br>Peak frequency (Hz) | Ctrl IgG CA1 -60 mV: 2 ± 1.2548 (n=14; N=3)<br>GBR IgG CA1 -60 mV: 1.9464 ± 1.4950 (n=14; N=5) | Mann-Whitney U-test, two-tailed | P = 0.8888<br>Z = -0.1398<br>U = 94.5 |
| | Ctrl IgG CA3 -60 mV: 0.1250 ± 0 (n=7; N=3)<br>GBR IgG CA3 -60 mV: 0.2917 ± 0.4308 (n=12; N=3) | Mann-Whitney U-test, two-tailed | P = 0.1829<br>Z = -1.3319<br>U = 31.5 |
| **E4e (left)**<br>Impedance \|Z\| (GΩ) | Ctrl IgG CA1 -75 mV: 0.1426 ± 0.0190 (n=14; N=3)<br>GBR IgG CA1 -75 mV: 0.1448 ± 0.0325 (n=14; N=5) | Two sample t-test, two-tailed | P = 0.8296<br>t = -0.2174<br>df = 26 |
| | Ctrl IgG CA3 -75 mV: 0.1112 ± 0.0201 (n=7; N=3)<br>GBR IgG CA3 -75 mV: 0.1025 ± 0.0245 (n=12; N=3) | Two sample t-test, two-tailed | P = 0.4357<br>t = 0.7983<br>df = 17 |
| **E4e (middle)**<br>Impedance \|Z\| (GΩ) | Ctrl IgG CA1 -75 mV: 0.0554 ± 0.0161 (n=14; N=3)<br>GBR IgG CA1 -75 mV: 0.0546 ± 0.0144 (n=14; N=5) | Two sample t-test, two-tailed | P = 0.8857<br>t = 0.1452<br>df = 26 |
| | Ctrl IgG CA3 -75 mV: 0.0482 ± 0.0189 (n=7; N=3)<br>GBR IgG CA3 -75 mV: 0.0479 ± 0.0208 (n=12; N=3) | Two sample t-test, two-tailed | P = 0.9714<br>t = 0.0364<br>df = 17 |
| **E4e (right)**<br>Peak frequency (Hz) | Ctrl IgG CA1 -75 mV: 2.5714 ± 1.5323 (n=14; N=3)<br>GBR IgG CA1 -75 mV: 2.2679 ± 1.5833 (n=14; N=5) | Mann-Whitney U-test, two-tailed | P = 0.7105<br>Z = 0.3712<br>U = 106.5 |
| | Ctrl IgG CA3 -75 mV: 0.1964 ± 0.1220 (n=7; N=3)<br>GBR IgG CA3 -75 mV: 0.2292 ± 0.2491 (n=12; N=3) | Mann-Whitney U-test, two-tailed | P = 0.7669<br>Z = 0.2965<br>U = 45 |
